# Structural mechanisms for gating and ion selectivity of the human polyamine transporter ATP13A2

**DOI:** 10.1101/2021.05.26.445856

**Authors:** Jordan Tillinghast, Darren Bowser, Kenneth Pak Kin Lee

## Abstract

Mutations in *ATP13A2*, also known as *PARK9*, cause a rare monogenic form of juvenile onset Parkinson’s disease named Kufor-Rakeb syndrome and other neurodegenerative diseases. *ATP13A2* encodes a neuroprotective P5B P-type ATPase highly enriched in the brain that mediates selective import of spermine ions from lysosomes into the cytosol via an unknown mechanism. Here we present three structures of human ATP13A2 bound to an ATP analogue or to spermine in the presence of phosphomimetics determined by electron cryo-microscopy. ATP13A2 autophosphorylation opens a lysosome luminal gate to reveal a narrow lumen access channel that holds a spermine ion in its entrance. ATP13A2’s architecture establishes physical principles underlying selective polyamine transport and anticipates a “pump-channel” intermediate that could function as a counter-cation conduit to facilitate lysosome acidification. Our findings establish a firm foundation to understand ATP13A2 mutations associated with disease and bring us closer to realizing ATP13A2’s potential in neuroprotective therapy.

**Highlights:**

1. Structures of the Parkinson’s disease-associated polyamine transporter ATP13A2
2. Structures of three transport cycle intermediates reveal the gating mechanism
3. Architecture of the polyamine binding site reveals mechanisms for ion selectivity
4. The polyamine binding site’s location anticipates an ion channel-like mechanism

## Introduction

Polyamines are ubiquitous in biology (Michael, 2016). Regulation of cell proliferation, modulation of ion channel behavior and neuroprotection are among the numerous pleiotropic functions of polyamines (Clarkson et al., 2004; Pegg, 2016; Yang et al., 2017). In humans, disruptions in polyamine metabolism and transport have been associated with cancer, aging and neurodegenerative diseases (Madeo et al., 2018). Polyamine uptake in a mammalian system was first shown by Dykstra and Herbst in 1965 (Dykstra and Herbst, 1965). More than half a century later, ATP13A2 was compellingly demonstrated to be a human polyamine uptake transporter (van Veen et al., 2020). ATP13A2 is a member of the P5B family of P-type adenosine triphosphatases (P5B-ATPases) highly expressed in the brain (Palmgren and Nissen, 2011; Schultheis et al., 2004; Sorensen et al., 2018). Polyamine export from lysosomes mediated by ATP13A2 sets the resting cellular polyamine content and protects against mitochondrial oxidative stress (van Veen et al., 2020; Vrijsen et al., 2020); its deficit impairs lysosome function and promotes neuronal cell death (van Veen et al., 2020). Many loss-of-function mutations in the *ATP13A2* gene (also known as *PARK9*) have been causally linked to a large spectrum of neurodegenerative diseases including Kufor-Rakeb syndrome and early-onset Parkinson’s disease (Bras et al., 2012; Di Fonzo et al., 2007; Estrada-Cuzcano et al., 2017; Klein and Westenberger, 2012; Lin et al., 2008; Ramirez et al., 2006; Spataro et al., 2019). Enhancement of ATP13A2 function has been proposed as a neuroprotective therapeutic strategy in Parkinson’s disease (Dehay et al., 2012a; Dehay et al., 2012b; Park et al., 2015).

Under physiological conditions the natural polyamines: spermine (SPM), spermidine (SPD) and putrescine (PUT) exist as tetravalent, trivalent and divalent cations, respectively (Bergeron et al., 1995). ATP13A2 prefers spermine over spermidine though both are efficiently transported; it shows little activity toward putrescine and metal ions (van Veen et al., 2020). The observation that ATP13A2 is able to transport ions with variable charge and size is surprising since canonical metal ion P-type ATPases including sarcoendoplasmic reticulum Ca^2+^-ATPase (SERCA) and Na^+^/K^+^-ATPase are highly selective ion pumps (Dyla et al., 2020; Morth et al., 2011). Given that acidic residues coordinating metal ions in SERCA and Na^+^/K^+^-ATPase are not conserved in ATP13A2 (Figure S1), we should expect to discover that ATP13A2 has evolved to handle polyamine ions using an alternate ion selection apparatus.

In addition to the question of ion selectivity, many fundamental aspects of polyamine transport by ATP13A2 remain to be characterized. What is the ion permeation pathway? What are the gating structures and how do they open and close? In the present study we address these questions by determining structures of human ATP13A2 representing three intermediates in its transport cycle using single-particle electron cryo-microscopy (cryo-EM). A biochemical analysis of structure-activity relationships among ATP13A2’s polyamine substrates is also presented.

## Results

### Biochemical characterization

We expressed full-length human ATP13A2 (hATP13A2) in a human embryonic kidney cell line and purified the recombinant ion pump in a detergent system containing dodecylmaltoside (DDM) and cholesteryl hemisuccinate (CHS) (Figures S2A and S2B). Purified hATP13A2 exhibited polyamine-dependent ATPase activity, in qualitative agreement with previous observations (Figures 1A and S2C) (van Veen et al., 2020). hATP13A2 is selective toward linear polyamines as the cyclic polyamine 1,4,8,11-tetraazacyclotetradecane (cyclam) did not stimulate hATP13A2 ATPase activity (Figures 1A and S2E).

**Figure 1.**
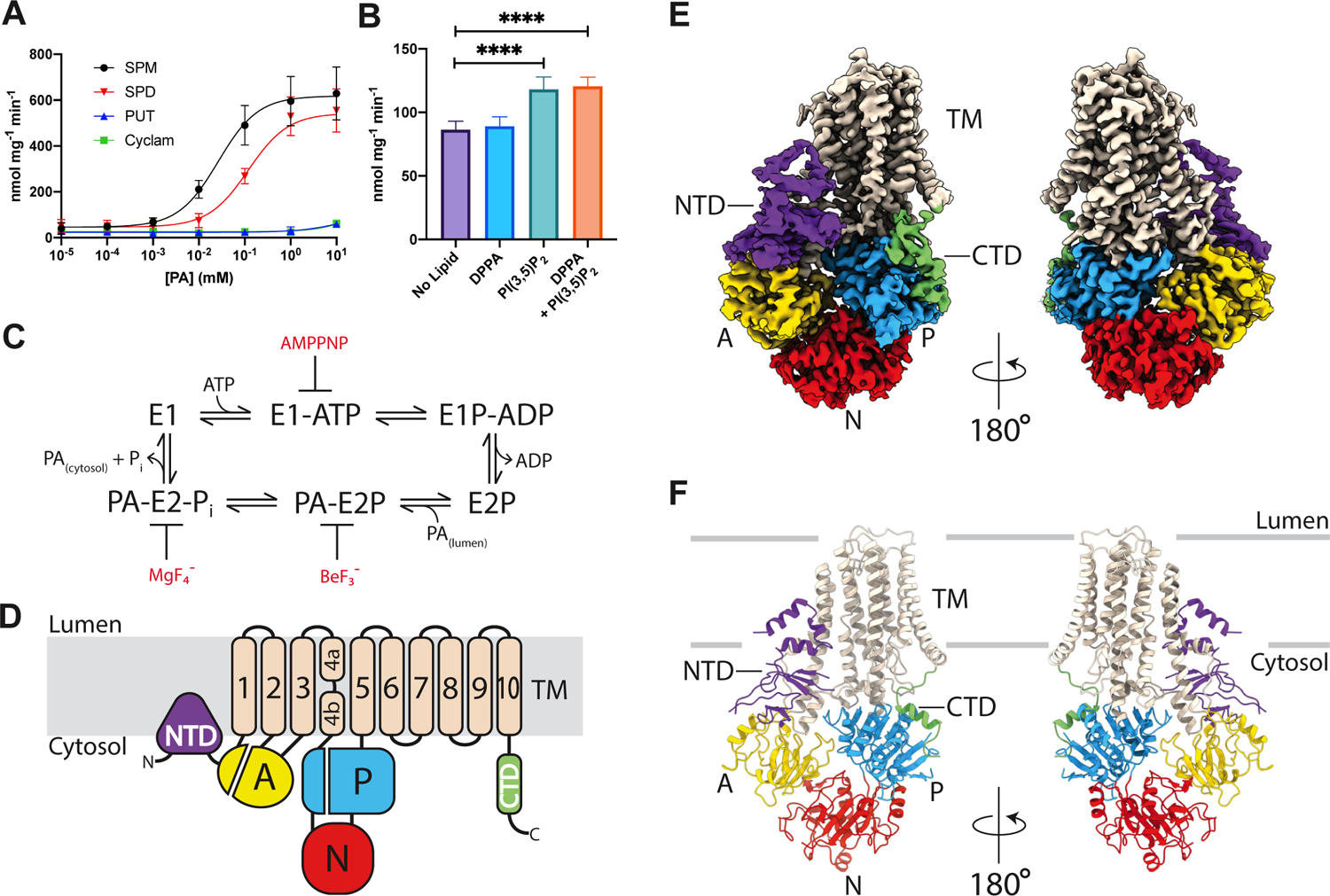
Architecture of human ATP13A2. (A) Polyamine-dependent ATPase activity of hATP13A2. Purified hATP13A2 exhibits dose-dependent polyamine stimulated ATPase activity characterized by apparent Michaelis constants: SPM (K_M,SPM_=26.4 µM), SPD (K_M,SPD_=118.3 µM). Putrescine and the cyclic polyamine 1,4,8,11-tetraazacyclotetradecane (cyclam) did not appreciably stimulate hATP13A2 ATPase activity and are assumed to have a K_M_>10 mM. 1 mM ATP was used in the reactions. Data points represent mean ± SEM (n=3). SPM – spermine, SPD – spermidine, PUT – putrescine. (B) Lipid dependence of hATP13A2 ATPase activity. ATPase activity of hATP13A2 in the presence of 1 mM SPM and 370 µM ATP with lipid supplementation. DPPA - dipalmitoylphosphatidic acid, PI(3,5)P_2_ –dipalmitoyl-phosphatidylinositol-3,5-bisphosphate. Data points represent mean ± SEM (n=3, ****P<0.0001). One-way ANOVA analysis was performed. (C) The reaction cycle of ATP13A2. Polyamine import takes place during the E2P to E1 half-cycle. Compounds used for stabilizing reaction intermediates are colored in red. PA – polyamine, P_i_ –inorganic phosphate. (D) Topology diagram of hATP13A2. Transmembrane helices M1 to M10 are numbered. TM – transmembrane domain, A – actuator domain, P – phosphorylation domain, N – nucleotide binding domain, NTD – N-terminal extension domain, CTD – C-terminal extension domain. The same domain abbreviations are used in all figures. (E) Cryo-EM density map of the E1-AMPPNP structure. (F) Ribbons representation of the atomic model of the E1-AMPPNP structure. The NTD contains a distinctive triangular “spade-like” structure containing three helices penetrating the inner-leaflet of the membrane (see also Figures S5A and S5B). This “spade” is cemented to the A domain via a *β*-sandwich structure. The CTD is composed by a linker that connects the end of M10 to an *α*-helix (*α*CTD) docked onto the surface of the P domain via hydrophobic contacts. Coloring as in (D).

Spermine-dependent ATPase activity of ATP13A2 is thought to be up-regulated by phosphatidic acid (PA) and phosphatidylinositol-(3,5)-bisphosphate (PI(3,5)P_2_) (van Veen et al., 2020). We found that spermine robustly stimulated hATP13A2 ATPase activity without necessitating lipid supplementation (Figures 1A and S2C); PI(3,5)P_2_ enhanced hATP13A2’s catalytic output but PA did not (Figure 1B). We attribute the differential effects of PA between our study and that of van Veen et. al. to differences in protein source and purification methods (van Veen et al., 2020).

### Structure determination and overall architecture

According to Post (Post et al., 1969) and Albers (Albers, 1967), ion transport by P-type ATPases can be understood as alternating cytosol-pump and extracellular milieu-pump ion exchange reactions mediated by four intermediates cycling in sequence: E1→E1P→E2P→E2 (Figure 1C). Intermediate states of hATP13A2 can be stabilized using the ATP analogue adenosine 5’-(*β*,*γ*-imido)-triphosphate (AMPPNP) and the phosphate analogues beryllium fluoride (BeF_3_^−^) and magnesium fluoride (MgF_4_^2−^). These inhibitors efficiently suppressed SPM-dependent hATP13A2 ATPase activity as expected (Figure S2D). We performed single-particle cryo-EM analysis of hATP13A2 complexed with: (i) AMPPNP (E1-AMPPNP); (ii) SPM plus BeF_3_^−^ (SPM-E2-BeF_3_^−^) and (iii) SPM plus MgF_4_^2−^ (SPM-E2-MgF_4_^2−^) (Figures 1C and S3A–S3F). The E1-AMPPNP structure represents ATP bound E1 (E1-ATP). The SPM-E2-BeF_3_^-^ structure represents the substrate bound phosphoenzyme (SPM-E2P). The SPM-E2-MgF_4_^2–^ structure represents the substrate bound dephosphorylated pump before PO_4_^3–^ is ejected (SPM-E2-P_i_). The three structures: E1-AMPPNP, SPM-E2-BeF_3_^−^ and SPM-E2-MgF_4_^2−^ were determined to resolutions of 2.9 Å, 2.7 Å and 3.0 Å, respectively (Figures S4E–S4G and Table S1).

A large transmembrane domain and a large multi-domain cytosolic headpiece comprise the hATP13A2 structures (Figures 1D, 1E, and S3D–S3F). The transmembrane regions of all three structures were similar in quality (Figures 1E, 3A, and S4H–S4J). The headpiece is most clearly resolved in the E1-AMPPNP structure, whose cryo-EM density was of excellent quality over most parts of the structure (Figures 1E and S4H). An atomic model of 1046 of 1180 residues in the hATP13A2 polypeptide sequence was built into the E1-AMPPNP cryo-EM map *de novo* (Figures 1E and 1F), which was then used as a template to build the SPM-E2-BeF_3_^−^ and SPM-E2-MgF_4_^2−^ structures.

A fold typical of P-type ATPases constitutes the core of hATP13A2 (Figure 1F). This core consists of ten transmembrane helices M1 through M10 and three cytosolic domains named: actuator (A), nucleotide binding (N) and phosphorylation (P) that make up the tripartite headpiece (Figures 1D–1F). The hATP13A2 core is flanked by two extensions: the N-terminal domain (NTD, aa. 1 to 186) embedded in the membrane inner leaflet (Figures 1D, S5A and S5B) and the C-terminal domain (CTD, aa. 1151 to 1180) (Figures 1D and 1F). Strong conservation of sequence indicates that both the NTD and CTD are structural signatures throughout the P5B-ATPase family (Figure S1).

### Functional implications of ATP13A2 extension domains

ATP13A2 is known to readily undergo autophosphorylation (Holemans et al., 2015; van Veen et al., 2020). This is not surprising since this step is expected to license ATP13A2 to sample “outward-open” conformations to access the lysosome lumen. How can the propensity of ATP13A2 to phosphorylate itself be explained from a physical perspective? In the autophosphorylated E2P state SERCA’s A domain positions Glu183 in the conserved TGES loop near the phosphorylated Asp351 sidechain in the P domain to catalyze Asp351’s dephosphorylation (Figure S1) (Olesen et al., 2007). The hATP13A2 E1-AMPPNP structure shows that the NTD anchors and pulls the A domain toward the inner leaflet of the lipid bilayer (Figure 2A). The resulting effect is the placement of the TGES loop in the hATP13A2 A domain ∼200% closer to the phosphorylated aspartic acid residue in the P domain (Asp513) compared to the analogous distance in the SERCA E1-AMPPCP structure (Figure 2A). The predisposition of ATP13A2 to autophosphorylate under physiological ATP settings can thus be understood simply as a reflection of a forward bias imposed by the NTD to attain the E2P conformation.

**Figure 2.**
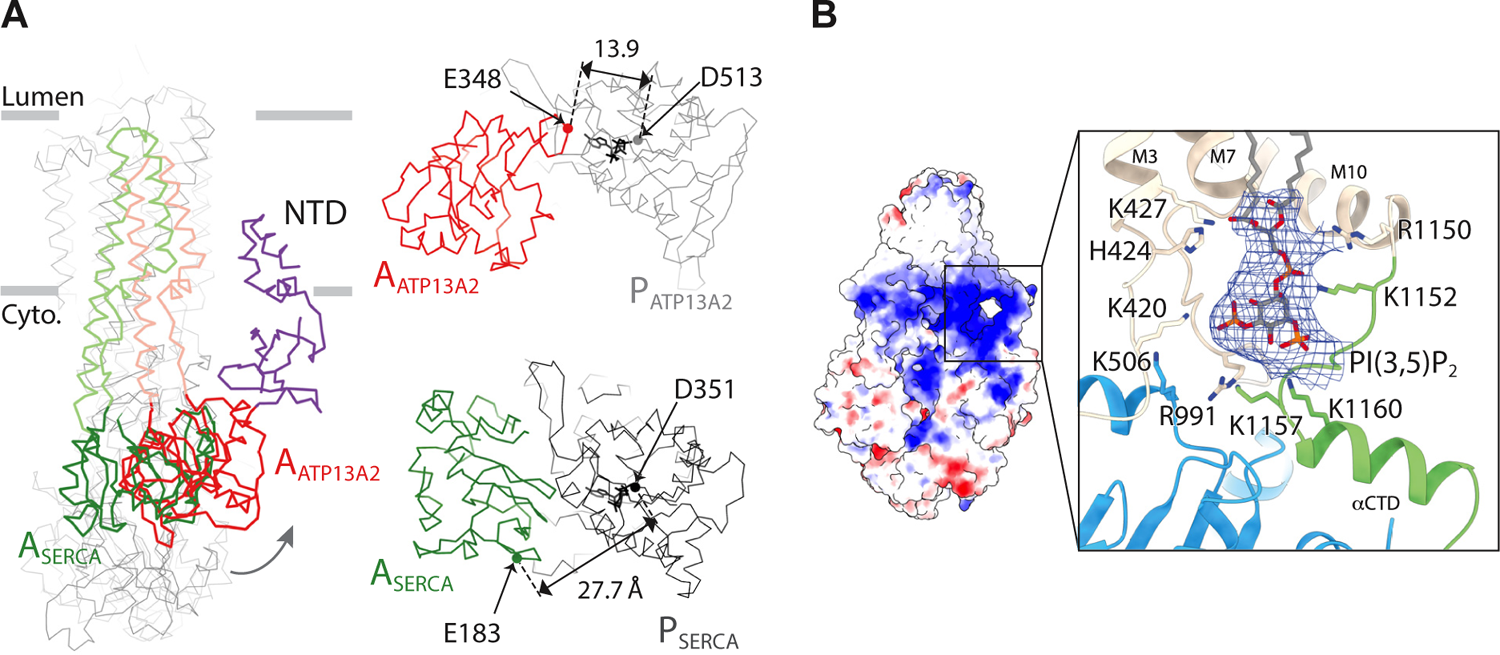
Function of ATP13A2 extensions. (A) Comparison of hATP13A2 and SERCA structures. The hATP13A2 E1-AMPPNP and SERCA E1-AMPPCP (PDB ID 3N8G) structures were aligned over their M7-M10 supporting domains and shown as grey and black *α*-carbon traces, respectively. A domains and the attached M1-M2 segments of hATP13A2 and SERCA are colored red and green, respectively. The hATP13A2 NTD is colored purple. Sideview (left - superposition) and top views (top-right – hATP13A2 and bottom-right – SERCA) are shown. Catalytic glutamate in the TGES motif and phosphorylated aspartate residues in the P domain are indicated by solid circles. The NTD lowers the energy barrier toward formation of the “outward-open” E2P conformation by bringing the A domain closer to its destination in the reaction coordinate. Cyto. – cytosol. (B) Putative PI(3,5)P_2_ binding site. Electrostatic potential surface of the hATP13A2 E1-AMPPNP structure (left - sideview). Close-up view of the putative PI(3,5)P_2_ binding site (boxed region on the left) shown in ribbons presentation (right). Sidechains are shown in ball-and-stick representation. Cryo-EM density corresponding to the modelled PI(3,5)P_2_ is shown as a blue mesh. Coloring as in Figure 1F.

To identify the location of the PA/PI(3,5)P_2_ binding site we consider the surface electrostatics of hATP13A2, which shows an extensive positively charged pocket located at bilayer-water interface near the CTD (Figure 2B). Strong cryo-EM density resembling the headgroup and glycerol backbone of a bound phospholipid molecule is present in the cationic pocket (Figure 2B). The exact identity of the occupant phospholipid remains to be determined and it likely corresponds to a co-purified endogenous lipid molecule. We model the observed cryo-EM density as PI(3,5)P_2_ on the basis of hATP13A2’s robust spermine-dependent ATPase activity and its further potentiation by PI(3,5)P_2_ (Figures 1A and 1B). What is clear is that this putative PI(3,5)P_2_ binding site is strategically located to serve as a communication nexus capable of relaying its lipid occupancy status to the transmembrane and cytosolic domains of hATP13A2. More work will be needed to establish the exact mechanism of lipid regulation.

### Ion permeation pathway and gating mechanism

The transmembrane domain of the E1-AMPPNP structure occludes a small cavity near the center of the membrane we refer to as the “occlusion chamber” (Figures 3B and S5C). Inside this chamber, the polar sidechains of Tyr259, Tyr940 and Gln944 coordinate the Asp967 carboxyl-sidechain through H-bonds; this tetrad of interacting amino acids is invariant within the P5B-ATPase family (Figures 3B, S1, and S5C). The Asp967 sidechain is likely protonated in the occluded E1-AMPPNP structure since the converse scenario of burying a negatively charged carboxylate-group in the membrane would be energetically prohibitive without a paired counter-charge. We do not know whether counter-transport of protons into the lysosome lumen is involved in the hATP13A2 transport cycle at present. However, the occlusion of Asp967 suggests that it may function as a proton carrier in the E1→E2P transition.

**Figure 3.**
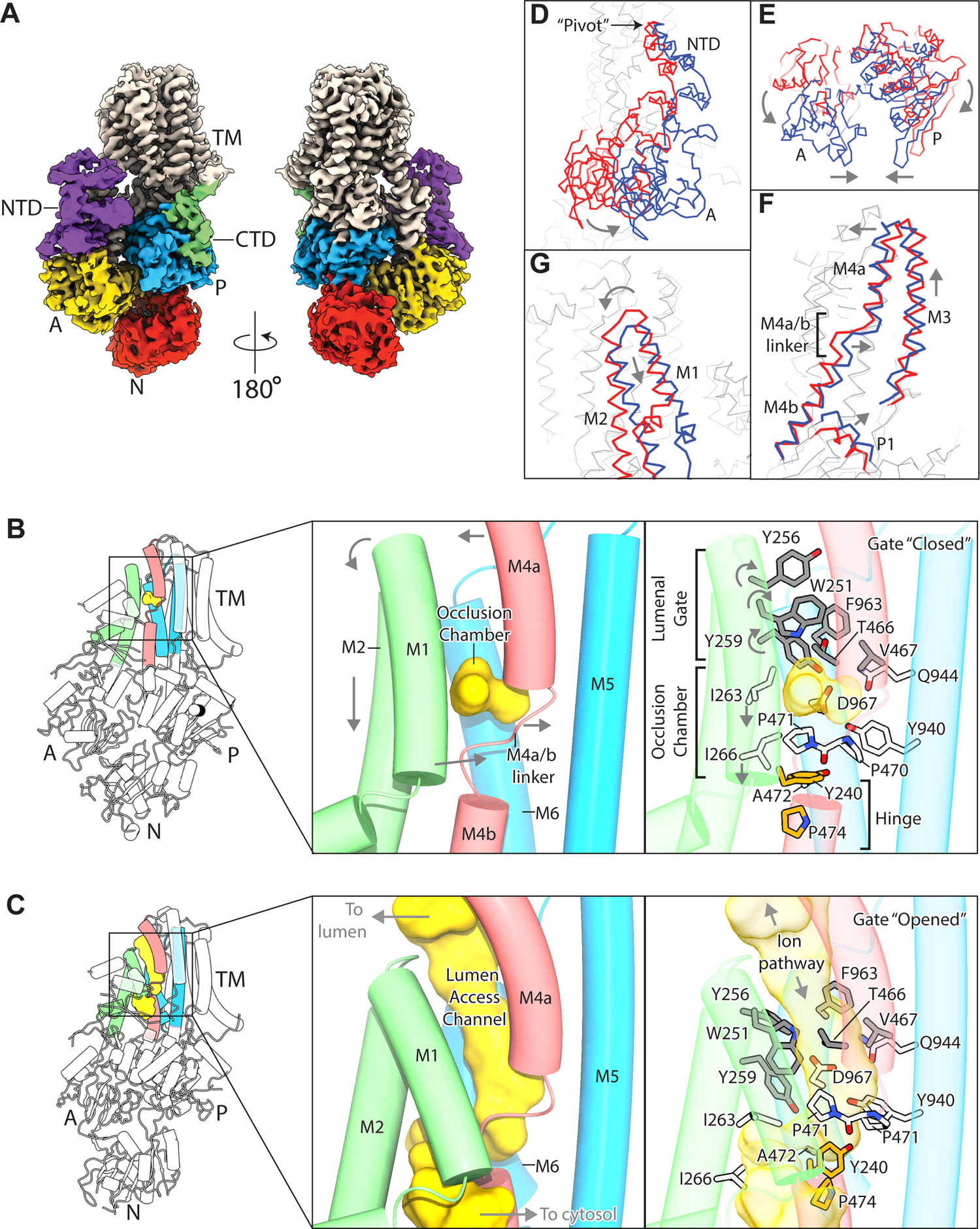
Gating mechanism of the lumen access channel. (A) Cryo-EM map of the SPM-E2-BeF3^−^ structure. Coloring as in Figure 1D. (B) The occlusion chamber. Cartoon representation of the E1-AMPPNP structure with helices shown as cylinders. M1-M2, M4a-M4b, and M5-M6 are colored green, red and blue, respectively. Sidechains of residues corresponding to the luminal gate, occlusion chamber and hinge are colored gray, white and orange, respectively. The volume enclosed by the occlusion chamber is shown as a yellow surface. Conformational changes associated with autophosphorylation and ADP dissociation are indicated by arrows. Polar and hydrophobic residues from M1, M2, M5 and M6 line the walls of the occlusion chamber and the conserved “PPAL” motif in the M4a/b linker forms the base. The luminal gate is “closed” in the E1-AMPPNP structure. (C) The lumen access channel. Cartoon representation of the SPM-E2-BeF_3_^−^ structure with helices shown as cylinders. The lumen access channel is shown as a yellow surface. Coloring as in (B). Contact between the base of M1 and top of M4b is maintained during the coupled movement of M1 and M2 through the interaction between Tyr240 and Pro474, which serves as a hinge for the motion. The displacement of M1 and M2 induced by autophosphorylation and ADP dissociation enables the aromatic sidechains of residues Trp251, Tyr256 and Tyr259 to rotate away from Thr466, Val467 and Phe963; this dissolves the hydrophobic plug and “opens” the luminal gate, revealing the lumen access channel. (D to G) Conformational changes coupled to autophosphorylation and ADP dissociation. E1-AMPPNP and SPM-E2-BeF_3_^−^ structures were aligned over their M7-M10 supporting domains and shown as red and blue *α*-carbon traces, respectively. Arrows are used to indicate direction of motion. (D) The A domain swings forward to displace the N domain from the P domain following Asp513 phosphorylation and ADP release. The trajectory of the A domain’s swing is constrained by its association with the NTD whose “spade” acts as a pivot for this motion. (E) The A domain and the P domain also rotate toward each other, bringing the TGES motif near Asp513 to catalyze dephosphorylation of the latter in a subsequent step. (F) Rotation of the P domain moves its P1 helix up toward the membrane and exerting a force on the cytosolic end of M4b. This causes the M4a-M4b segment to buckle at the M4a/b linker. (G) The swinging motion of the A domain exerts a pull on M1 and M2 through direct attachments resulting in the rotation and downward translation of these transmembrane segments toward the cytosol.

Access to the occlusion chamber from the lysosome lumen is controlled by an ∼13 Å hydrophobic barrier formed by M1, M2, M4a and M6 we call the “luminal gate” (Figure 3B). In the E1-AMPPNP structure the residues Trp251, Tyr256, Tyr259, Thr466, Val467 and Phe963 converge to form a hydrophobic plug “closing” the luminal gate (Figure 3B). In the SPM-E2-BeF_3_^−^ structure the luminal gate “opens” to reveal a narrow “lumen access channel” penetrating 15 Å into the membrane to join the lumen and occlusion chamber (Figures 3B and 3C). The narrow architecture of this access tunnel (∼3.2 Å in diameter) departs strikingly from the wide polar funnel (15 to 4 Å in diameter) observed in the E2P structure of SERCA that connects the ER lumen and Ca^2+^ binding sites (Olesen et al., 2007). A comparison of the E1-AMPPNP and SPM-E2-BeF_3_^−^ structures reveals an allosteric mechanism by which autophosphorylation of Asp513, mimicked by BeF_3_^−^ binding, in the P domain opens the luminal gate 59 Å away (Figures 3D–3G and Video S1, refers to Figure 3).

Structural alignment of the SPM-E2-BeF_3_^−^ and SPM-E2-MgF_4_^2−^ structures shows that they are very similar andare both in the outward-facing conformation (Figures S3G–S3I, root mean square deviation=0.52 Å). This is in strong contrast to SERCA, which undergoes large conformational changes from an outward-facing state to an occluded state in the E2P→E2-P_i_ reaction (Olesen et al., 2007). The important implication here is that structural transitions bringing ATP13A from the spermine bound outward-facing state to the next major conformation in the transport cycle must occur after PO_4_^3-^ dissociation in the E2-P_i_→E1 step (Figure 1C). This next transport cycle intermediate may well be an occluded conformation though its exact nature remains to be determined. These observations underscore the diversity in coupling mechanisms between enzymatic and gating transformations among all P-type ATPases, our understanding of which is still clearly incomplete.

### Structural basis of spermine recognition

Electrostatic potential surface analysis of the SPM-E2-BeF_3_^−^ structure shows that the lumen access channel possesses a strongly negative charge character, consistent with a possible role in binding positively charged polyamines (Figure 4A). A cigar-shaped cryo-EM density extends from the entrance of the lumen access channel 11 Å down to the occlusion chamber (Figure 4B). Spermine can be easily modelled into this density such that the spermine amine groups come into direct contact with acidic residues in this site (Figure 4C). Therefore, the outer portion of the lumen access channel is the polyamine binding site. In the following discussion we will refer to the four spermine amine groups as N1, N5, N10 and N14, respectively since spermine is a linear chain of 14 non-hydrogen atoms (Figures 4D, 4E, and S2E).

**Figure 4.**
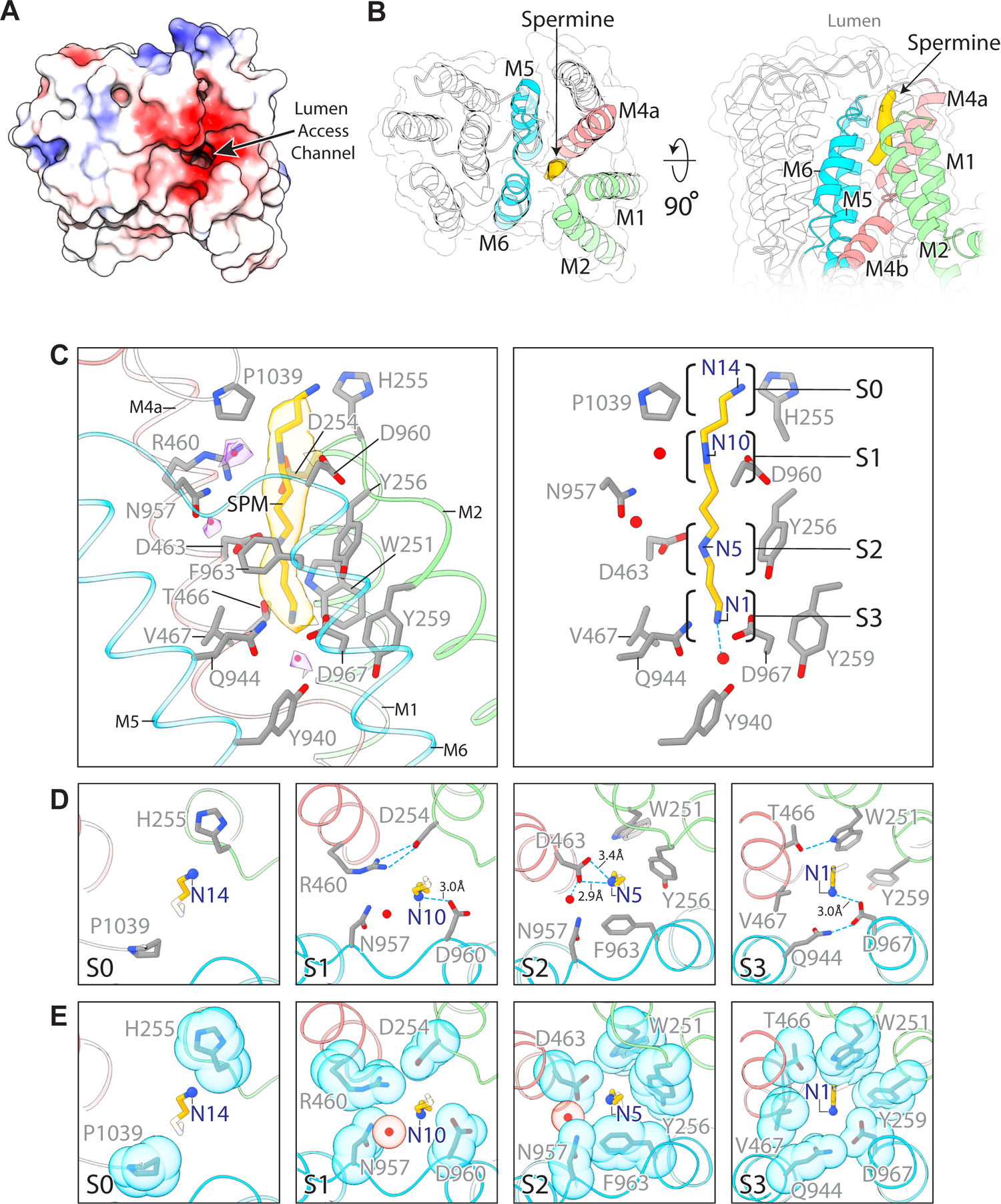
Structural basis of spermine recognition. (A) Electrostatic potential surface of the lumen access channel entrance viewed from the lumen. (B) Location of the polyamine binding site. The transmembrane domain of hATP13A2 is shown in ribbons representation. Coloring of helices as in Figure 3C. Topview (left) is in the same perspective as in (A). Sideview is shown on the right. Cryo-EM density of the bound spermine molecule is colored yellow. (C) Architecture of the polyamine binding site. Close-up view of the polyamine binding site (left). Spermine and residues lining the polyamine binding site are colored yellow and gray, respectively and shown in ball-and-stick representation. Helices colored as in (B). Cryo-EM densities of the bound spermine and solvent are shown as transparent surfaces colored yellow and purple, respectively. Water is shown as red spheres. Amine group binding sites S0 to S3 are indicated by brackets (right). Some polyamine binding site residues are omitted in the right panel for clarity. (D and E) Interactions between spermine amine groups and the polyamine binding site. Topviews of S0 to S3 are shown. Coloring as in (C). Transparent van der Waals surfaces of sidechains and solvent molecules in S0 to S3 are shown in (E). His255 and Pro1039 make up S0 and do not come into close contact with N14. As a result, N14 is less ordered and is free to interact with solvent molecules. In S1, N10 interacts directly with Asp960 through coulombic interactions while a water molecule associated with Asn957 and an Asp254-Arg460 salt-bridge surround N10 in a polar environment. The chemical environment of S2 is very different from S1 and is the narrowest part of the polyamine binding site (diameter≈3.2 Å, Figure 6C). In S2, a completely dehydrated N5 interacts with both oxygen atoms of the Asp463 sidechain carboxyl group while Trp251, Tyr256 and Phe963 encircle N5 on three sides with aromatic sidechains. In S3, N1 interacts tightly with Asp967 and is further surrounded by a combination of polar and hydrophobic residues: Trp251, Tyr259, Thr466, Val467 and Gln944. Additionally, N1 interacts with a water molecule kept just below S3.

The polyamine binding site overlaps with the luminal gate and occlusion chamber and comprises residues from the external portion of M1, M2, M4a, M5 and M6 (Figures 4B and 4C). Spermine is held in an almost fully extended conformation at the polyamine binding site and is partially dehydrated. N1, N5 and N10 are buried inside the lumen access channel while N14 is exposed to solvent (Figures 4C–4E). For simplicity we will refer to the four sites occupied by N14, N10, N5 and N1 as: S0, S1, S2 and S3, respectively (Figure 4C). Since the polyamine binding site is both narrow and embedded halfway inside the electric field of the membrane, the binding of polyamines to hATP13A2 is likely voltage dependent. Based on conservation of 8 of 14 residues in the polyamine binding site (Figure S1), we propose that other members of the P5B-ATPase family are also polyamine transporters. Consistent with this notion, the P5B-ATPase ATP13A3 was found to mediate polyamine transport (Hamouda et al., 2020).

### Ion dehydration and selectivity

A fundamental challenge all selective ion transport systems must overcome is the requirement for hydrated ions in solution to be transferred to their binding sites in a completely or partially dehydrated state (Hille, 2001). Ion dehydration incurs significant energetic cost that must be offset by compensating interactions (Gouaux and Mackinnon, 2005). We can understand how hATP13A2 solves the dehydration problem by considering the manner of spermine binding in the polyamine binding site.

The narrow channel architecture of the polyamine binding site constrains N1 to diffuse through S0→S1→S2→S3 in sequence during the spermine binding reaction (Figures 4C–4E). The structure of the polyamine binding site tells us that it is designed to dehydrate polyamine ions in a stepwise fashion: spermine becomes progressively more dehydrated as it migrates deeper into this narrow passage. The significance of this “multistep dehydration” mechanism is that it breaks down the dehydration process into a series of smaller steps with lower kinetic barriers, thus allowing polyamine binding to proceed in a more facile way.

Multistep dehydration can be considered a selectivity mechanism against metal ions; we would expect this effect to favor binding of linear polyamines and not metal ions. hATP13A2 utilizes an additional mechanism of metal ion counter-selection. In selective metal ion channels and transporters, including SERCA, dehydrated metal ions are chelated by a constellation of five to eight oxygen atoms that replaces the inner hydration sphere (Figure S6C) (Gouaux and Mackinnon, 2005; Sorensen et al., 2004; Toyoshima et al., 2000; Yamashita et al., 2005; Zhou et al., 2001). In hATP13A2, the amine binding sites S1, S2 and S3 can each provide at most two oxygen atoms from carboxylate groups and/or water (Figures 4D–4E). This low coordination number, though apparently sufficient to stabilize dehydrated amine groups in polyamines, is less than that needed to stabilize naked metal ions. Thus, the hATP13A2 polyamine binding site is, by design, not built to handle dehydrated metal ions. We refer to this ion selectivity mechanism as “chelate deprivation”.

### The origin of spermine selectivity

How can we rationalize hATP13A2’s polyamine selectivity profile? Perhaps the most obvious intuition derived from inspecting the polyamine binding site is that hATP13A2 must prefer linear polyamines over cyclic polyamines, as is observed (Figures 1A and S2E). Cyclization constrains the latter to have a minimum cross-section area too large even for the polyamine binding site entrance. Linear polyamines are unconstrained polymer chains and can thus thread through the narrow binding site.

Since putrescine and spermidine are essentially fragments of spermine we can deduce by analogy how they might be docked in the polyamine binding site based on spermine’s binding configuration (pose I in Figures 5A and S2E). This analysis predicts that putrescine and spermidine can form two or three ionic interactions with the polyamine binding site, respectively. It follows that hATP13A2’s impressive ∼100-fold selectivity for spermidine over putrescine (Figure 1A) is explained in part by avidity effects stemming from a greater number of possible ion-pairs formed by spermidine. Avidity is a selectivity mechanism not available to metal ion transporters. Spermidine binding also buries a larger hydrophobic surface compared to putrescine; thus, stronger spermidine binding is explained additionally by the hydrophobic effect.

**Figure 5.**
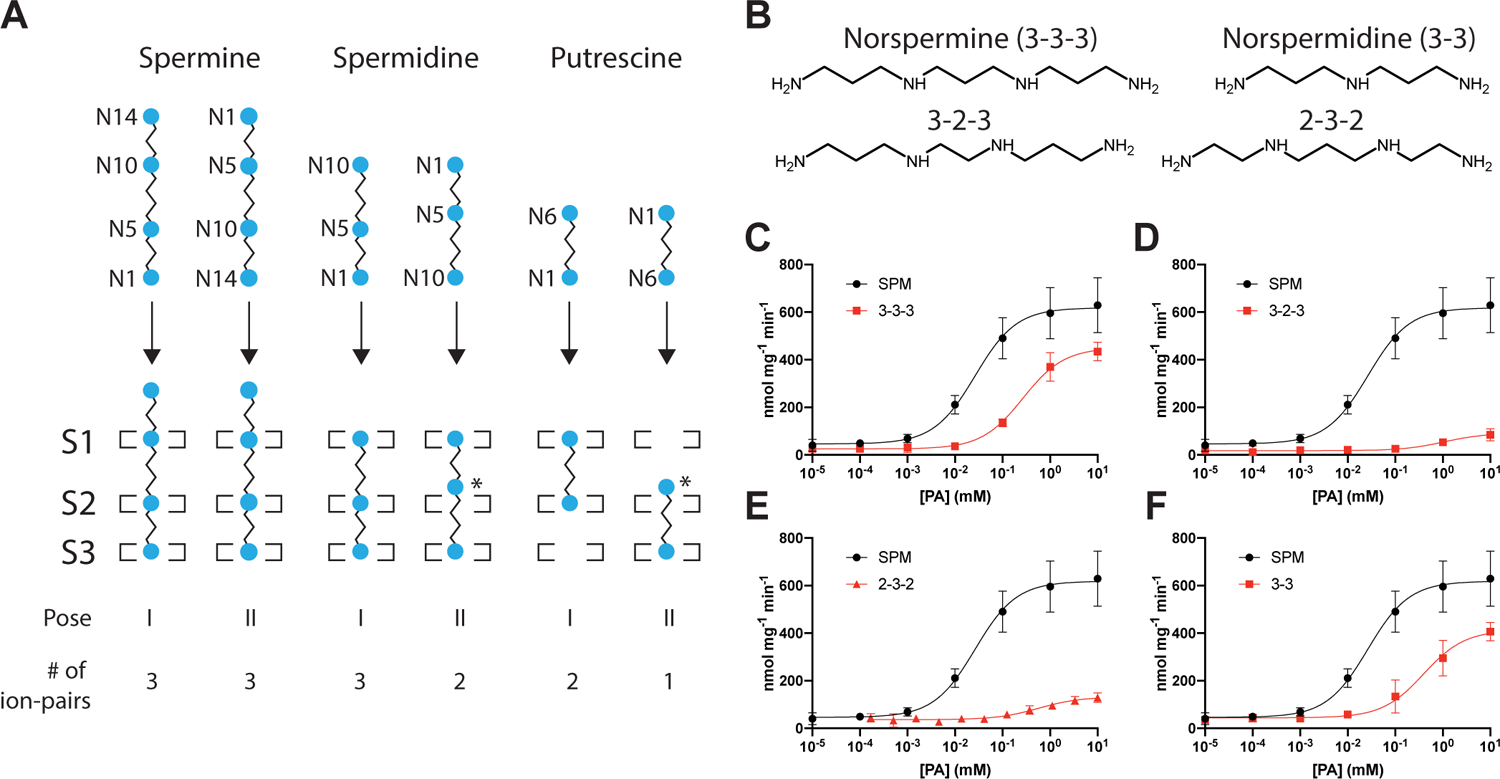
Origin of spermine selectivity. (A) Model of polyamine binding to hATP13A2. Amine and aliphatic groups are shown as blue circles and black sticks, respectively. Amine binding sites S1 to S3 are drawn as square brackets. Spermine, spermidine and putrescine can enter the polyamine binding site in two possible orientations (Pose I and II). Suboptimal interactions are indicated by asterisks. Spermine bound in pose I and pose II are chemically indistinguishable configurations due to its symmetry. In pose II spermidine makes what must be a weaker interaction with the polyamine binding site compared to pose I as only two of three possible ion-pairs can be formed. Putrescine can adopt both poses shown in any orientation. (B) Spermine and spermidine analogs used for structure activity relationship analysis. A shorthand denoting the length of aliphatic linkers between amine groups is used. 3-3-3 – norspermine, 3-2-3 – 1,2-Bis(3-aminopropylamino)ethane, 2-3-2 – N,N′-Bis(2-aminoethyl)-1,3-propanediamine, 3-3 – norspermidine. (C to F) Dose-response curves showing the effect of spermine and spermidine analogs shown in (B) on hATP13A2 ATPase activity in the presence of 1 mM ATP. Apparent Michaelis constants: norspermine (K_M,3-3-3_=267.0 µM), norspermidine (K_M,3-3_=394.5 µM). 3-2-3 and 2-3-2 did not appreciably stimulate ATPase activity. Abbreviations as in (B). Results from the spermine titration in Figure 1A shown as a reference. Error bars represent mean ± SEM (n=3).

hATP13A2 has a more modest ability to discriminate spermine from spermidine (Figure 1A). Binding of spermine and spermidine in the polyamine binding site would involve formation of the same three ion-pairs and buries the same area of hydrophobic surface in our model (Figure 5A). Thus, the contribution of avidity and hydrophobic effects to binding is the same for both spermine and spermidine. How can hATP13A2’s ∼4-fold spermine preference over spermidine be explained? To understand the origin of hATP13A2’s selectivity for spermine we consider the symmetry present in the chemical structure of spermine itself. Spermine and spermidine can enter the polyamine binding site in two possible orientations depicted in Figure 5A as pose I and pose II. What emerges from this picture is that spermine intrinsically has a higher probability of engaging the polyamine binding site in a high affinity configuration compared to spermidine. hATP13A2’s apparent preference for spermine over spermidine is thus, at least partly, a reflection of this underlying relationship (Figure 5A).

The selectivity mechanism above presupposes that the polyamine binding site must be sensitive to the degree of spatial matching between amine groups in the bound polyamine molecule and the amine binding sites S1, S2 and S3. The degree of spatial matching can be affected by binding orientation, as discussed above, and also by varying the spacing between amine groups in polyamines. To test the latter, we characterized the effect on ATPase function of a small but focused panel of spermine analogs with varying amine group spacings (Figures 5B–5F). Symmetry in spermine and its symmetrical analogs restricts hATP13A2 binding to a single possible configuration. We found that norspermine, in which a single methylene (-CH_2_-) group from the 4-carbon N5-N10 linker in spermine is removed, exhibits a 10-fold reduction in hATP13A2 binding affinity compared to spermine (Figure 5C and S2E). Further elimination of an additional -CH_2_-group from the N5-N10 linker or both N1-N5/N10-N14 linkers from norspermine almost completely abolished hATP13A2 ATPase stimulation in both cases (Figures 5D, 5E and S2E). Thus, the polyamine binding site has evolved to read the length of aliphatic spacers between amine groups in polyamines. The relative positions of S1-S2 and S2-S3 are tuned for recognition of 4-carbon and 3-carbon spacings between amine groups in polyamines, respectively.

The fact that norspermine can potentiate hATP13A2 ATPase function enables us to test the idea that symmetry accounts for spermine’s enhanced binding affinity compared to spermidine. If this idea is valid, then one would predict norspermidine to be poorly distinguished from norspermine as both polyamines would make the same interactions with hATP13A2 in all possible orientations due to their inherent symmetry (Figure 5B). This prediction is indeed borne out by our observations (Figures 5C and 5F). Therefore, molecular symmetry is a determinant of polyamine affinity for hATP13A2.

### Putative cytosolic ion exit pathway

At the base of the polyamide binding site where it meets the “PPAL” motif in the M4a/b linker the lumen access channel makes a sharp turn and continues toward a side portal between M2 and M6 facing the lipid bilayer (Figures 6A and 6B). The lumen access channel then terminates at a narrow constriction between M2 and M4b formed by Ala472, Ile263 and Ile266 that leads to a funnel-shaped vestibule connected to the cytosol between M1, M2 and M4b (Figures 3C, 6B, and 6C). Ala472, Ile263 and Ile266 likely constitute a “cytosolic gate” since it is the thinnest physical barrier separating the lumen access channel and the cytosolic vestibule in the SPM-E2-BeF_3_^−^ structure (Figure 6C). A chain of ordered solvent molecules connects N1 of the bound spermine molecule and the cytosolic gate, tracing a putative cytosolic ion exit pathway (Figures 6A and 6B).

**Figure 6.**
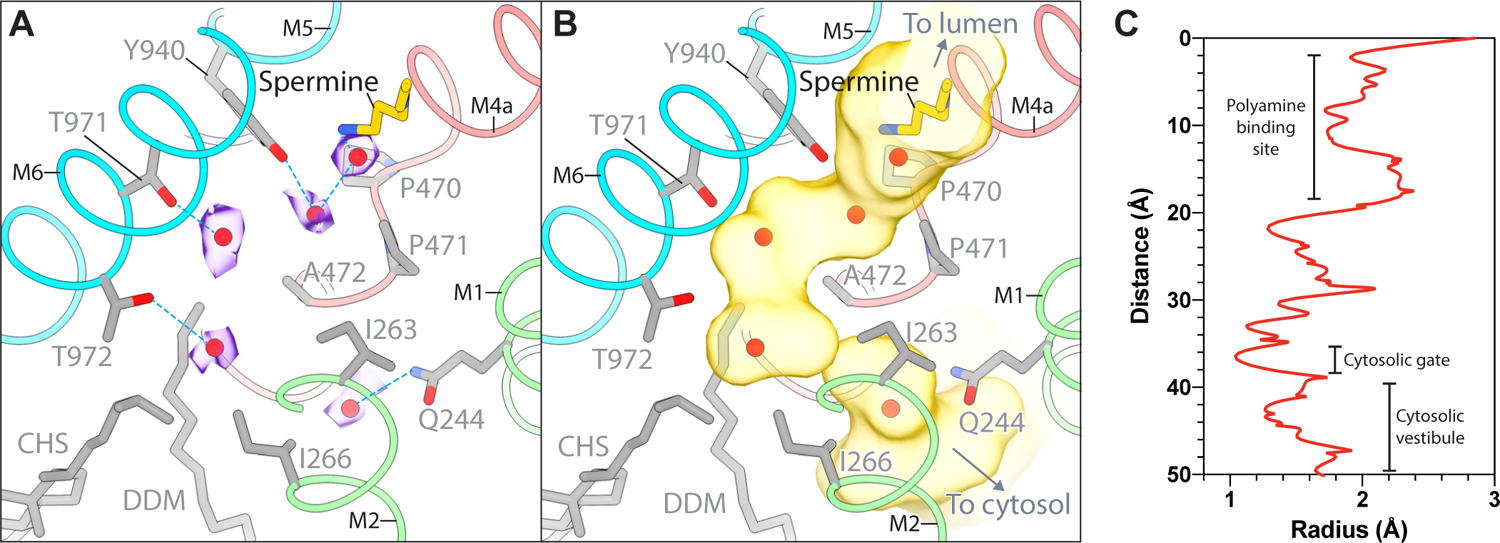
A putative cytosolic ion exit pathway. (A-B) A detailed view of the cytosolic gate region in the SPM-E2-BeF_3_^−^ structure is shown. The tunnel beneath the polyamine binding site leads to a narrow constriction referred to in the text as the “cytosolic gate” formed by Ile263, Ile266 and Ala472. On the other side of the cytosolic gate is a funnel-shaped “cytosolic vestibule” that leads to the cytoplasm. Ordered solvent molecules are shown as red spheres. Cryo-EM density corresponding to solvent is shown as transparent purple surfaces in (A). The lumen access channel and the cytosolic vestibule are rendered as transparent yellow surfaces in (B). Helices are colored as in Figure 3C. (C) Tunnel radius along the ion pathway. A plot of the tunnel radius of the lumen access channel and cytosolic vestibule estimated using MOLE 2.5 is shown. The locations of the polyamine binding site, cytosolic gate and cytosolic vestibule are indicated.

## Discussion

P-type metal ion pumps including SERCA and Na^+^/K^+^-ATPase have internal ion binding sites flanked by gates embedded near the center of the lipid membrane consistent with an alternating access transport mechanism (Drew and Boudker, 2016; Dyla et al., 2020; Jardetzky, 1966). The ATP13A2 polyamine binding site’s location in the external portion of the lumen access channel is therefore surprising. Could hATP13A2 residues in positions analogous to Ca^2+^ binding sites in SERCA form an internal binding site that occludes spermine or other polyamines in a subsequent step? This scenario is unlikely, in part because structural alignment of SERCA and hATP13A2 transmembrane domains shows that residues involved in Ca^2+^ coordination in SERCA are not conserved in hATP13A2 (Figure S6). We did not find additional cavities in the transmembrane domain that might potentially serve as polyamine occlusion sites.

Overlap between the luminal gate and polyamine binding site implies that a bound polyamine molecule will act as a wedge preventing closure of the luminal gate. This is incompatible with an alternating access transport mechanism as the latter involves ion occlusion, which cannot happen if the luminal gate is unable to close (Drew and Boudker, 2016; Jardetzky, 1966). The external location of the polyamine binding site, and the single-file arrangement of mobile charges within it, are actually features reminiscent of the selectivity filter of ion channels (Zhou et al., 2001). The selectivity filter is the narrowest part of an ion conducting pore optimized for selectivity and throughput (Gouaux and Mackinnon, 2005; Hille, 2001). We propose that hATP13A2 may sample an ion channel-like intermediate state — a “pump-channel” in which both the luminal gate and cytosolic gate are open following PO_4_^3−^ release from the polyamine bound E2-P_i_ state (Figure 7). In this intermediate state the polyamine binding site acts as a selectivity filter to enforce selective ion movement across the membrane. This mechanism could explain selective polyamine transport by hATP13A2 without invoking the existence of an ion occlusion chamber. We know at least one other P-type pump that can function as an ion channel: the palytoxin-modified Na^+^/K^+^-ATPase pump-channel described by Artigas and Gadsby (Artigas and Gadsby, 2002). Perhaps ATP13A2 evolved to exploit an ion channel-like state during its transport cycle, independent of toxin binding. This pump-channel model should bear consideration because it suggests the tantalizing possibility that hATP13A2 may act as a spermine/spermidine conduit to depolarize the lysosome and thus promote V-type ATPase driven proton flux (Mindell, 2012). Such a function could account for the impaired lysosome acidification observed in the settings of Parkinson’s disease-linked ATP13A2 mutations and targeted ATP13A2 knock-out (Dehay et al., 2012b; van Veen et al., 2020). The results presented here offer fertile ground to understand more deeply the mechanisms and implications of ATP13A2-mediated transport and regulation in the context of lysosome function and neurodegeneration.

**Figure 7.**
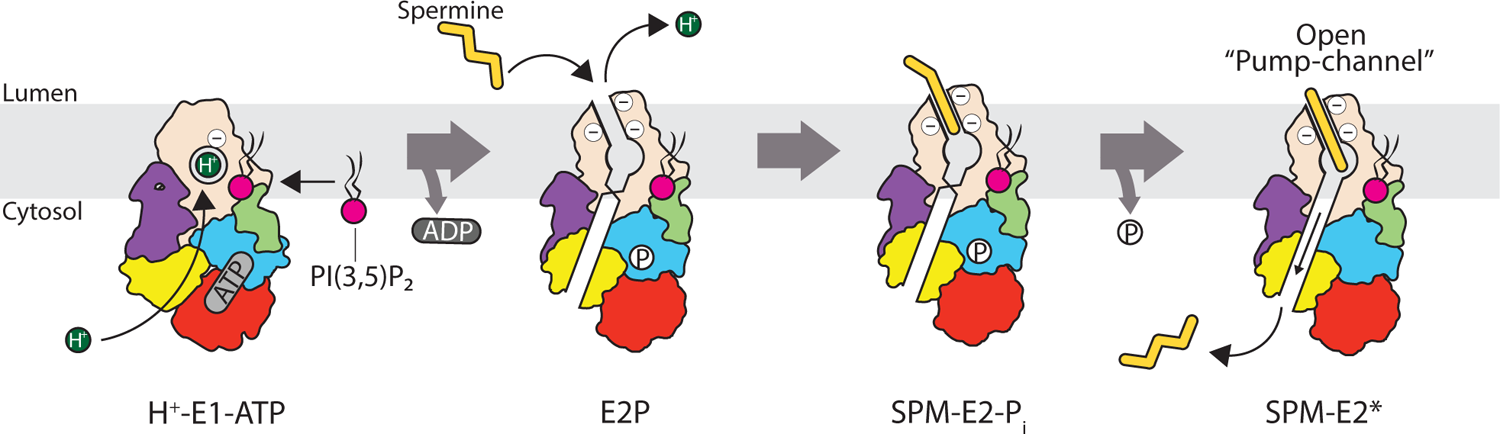
Proposed mechanism of polyamine transport. Since Asp967 is likely uncharged when it is occluded, we suggest that it obtains a proton from the cytosol during the E1→E1-ATP step. Autophosphorylation and ADP release precedes the E2P state, at which point polyamine and/or proton exchange with the lysosomal lumen takes place. Binding of spermine or spermidine stimulates ATP13A2 dephosphorylation. Release of PO_4_^3−^ from the ATPase active site triggers formation of a “pump-channel” intermediate, in which both luminal and cytosolic gates are open, thus allowing polyamine translocation into the cytosol. Binding of PI(3,5)P_2_ or PA to the PI(3,5)P_2_ binding site potentiates the transport cycle.

The new ATP13A2 structures permit a confident mapping of disease mutations onto two distinct conformational states in the transport cycle of ATP13A2 (Figure S7). Mutations associated with hereditary spastic paraplegia and neuronal ceroid lipofuscinosis are concentrated near the catalytic site in the P-domain and likely impact ATP binding and/or autophosphorylation. The remaining mutations associated with Kufor-Rakeb syndrome and early-onset Parkinson’s disease are distributed throughout the ATP13A2 structure and possibly affect the folding, maturation and/or stability of the protein. The precise explanations as to why particular mutations are associated with a specific disease phenotype will require additional study. Notably, disease-associated loss-of-function mutations are conspicuously absent from the polyamine binding site and the ion permeation pathway. The underlying reasons for this curious pattern are unclear and may warrant further investigation. However, the ATP13A2 structures described here provide a firm foundation to begin rationalizing ATP13A2 mutations associated with neurodegenerative disorders.

In summary, this work describes cryo-EM structures of the Parkinson’s disease linked polyamine transporter ATP13A2 in occluded and spermine-bound states. Comparison of these structures reveal the ion permeation pathway and gating mechanism. Our structural and functional analysis establishes the physical principles of selective polyamine transport. These principles are relevant to any mechanistic consideration of other polyamine transport systems found in animals, bacteria, fungi and plants since the problem of transporting linear organic polycations across membranes is fundamentally the same in all organisms (Fujita et al., 2012; Igarashi and Kashiwagi, 1999). The surprising location of the polyamine binding site and its implications for the transport mechanism highlights the P-type ATPase fold as an impressively malleable platform for catalyzing the movement of polar substances across membranes. Structures and mechanisms presented here provide useful counterpoints to understand the operation of all P-type ATPases transporting disparate materials including metal ions, protons, lipids and peptides (Abe et al., 2018; Huang et al., 2017; McKenna et al., 2020; Morth et al., 2007; Olesen et al., 2007; Pedersen et al., 2007; Shinoda et al., 2009; Toyoshima et al., 2000).

## Conclusion

Our findings refine the current knowledge of ion transport mechanisms among P-type ATPases. The structures described here furnish rich opportunities for pharmacological investigation of ATP13A2s’s therapeutic potential in disease contexts including cancer and Parkinson’s disease (Madeo et al., 2018).

## Acknowledgments

We thank Mohamed Trebak, Christopher Yengo, Lisa Shantz and Donald Gill for critical reading of the manuscript and Chen Xu for help with SerialEM. This work was supported by The Pennsylvania State University start-up funds to K.L.

## Author contributions

J.T. and D.B. performed protein expression, purification and biochemistry experiments. D.B. performed the molecular biology experiments. K.L performed EM experiments. K.L. conceived and supervised the project. J.T. and K.L. analyzed the results. K.L. wrote the manuscript.

## Competing interests

Authors declare that they have no competing interests.

## Data and materials availability

Cryo-EM density maps of the hATP13A2 E1-AMPPNP, SPM-E2-BeF_3_^−^ and SPM-E2-MgF_4_^2−^ forms have been deposited in the electron microscopy data bank under accession code EMD-23683, EMD-23684 and EMD-23685, respectively. Atomic coordinates of the hATP13A2 E1-AMPPNP, SPM-E2-BeF_3_^−^ and SPM-E2-MgF_4_^2−^ have been deposited in the protein data bank under accession code 7M5V, 7M5X and 7M5Y, respectively. Materials are available from the corresponding author upon reasonable request.

## Methods

### Cell lines

Sf9 cells were cultured in Sf-900 II SFM medium (GIBCO) at 28°C. HEK293S GnTI^-^ cells cultured in Freestyle 293 medium (GIBCO) supplemented with 2% FBS at 37°C. Cell lines were acquired from and authenticated by American Type Culture Collection (ATCC). The cell lines were not tested for mycoplasma contamination.

### Protein expression and purification

cDNA encoding full-length human ATP13A2 isoform A (hATP13A2, UniProt ID Q9NQ11-1) was cloned into the pEG BacMam vector containing a C-terminal GFP preceded by a PreScission protease cleavage site (Goehring et al., 2014). hATP13A2 was expressed in HEK293S GnTI^−^ cells using the BacMam method. In brief, bacmids encoding the hATP13A2-GFP fusion were generated using DH10Bac cells according to the manufacturer’s instructions. BacMam baculoviruses were produced using *Spodoptera frugiperda* Sf9 cells cultured in SF900II SFM medium. For protein expression, suspension cultures of HEK293S GnTI^−^ cells cultured in Freestyle 293 medium were infected with BacMam baculovirus at a density of ∼1.5×10^6^ cells/ml. After 24 hrs at 37°C, the infected cultures were supplemented with 10 mM sodium butyrate and grown for a further 24 hrs at 30°C before harvesting. All subsequent manipulations were performed at 4°C.

For large-scale purification, cell pellets from 4 liters of culture were resuspended in lysis buffer containing 50mM Hepes, pH 7.5, 150mM NaCl, 2mM DTT and supplemented with a protease inhibitors cocktail (Lee et al., 2017). The cell suspension was disrupted in a Dounce homogenizer and the resulting lysate was clarified at 39800 x g for 30 min. The crude membrane pellet obtained was then resuspended once again in lysis buffer and was stirred for 2 hrs in the presence of 1.5% (w/v) lauryl maltose neopentyl glycol (LMNG) and 0.3% (w/v) cholesteryl hemisuccinate (CHS). The solubilized membranes were clarified by centrifugation at 39800 x g for 45 min and the resulting supernatant was mixed with GFP nanobody-coupled sepharose resin (prepared in-house) by rotation. After 60 min, the resin was collected and washed with 20 column volumes of wash buffer containing 20 mM Hepes, pH 7.5, 150mM NaCl, 1 mM DTT, 0.03% (w/v) dodecylmaltoside (DDM) and 0.0015% (w/v) cholesteryl hemisuccinate (CHS). PreScission protease was used to elute hATP13A2 from the GFP nanobody resin by overnight incubation. The eluted protein was concentrated to a volume of ∼500 uL prior to fractionation on a Superose 6 column equilibrated with 20 mM Hepes, pH 7.5, 150mM NaCl, 1 mM DTT, 0.03% (w/v) DDM and 0.0015% (w/v) CHS. Peak fractions were collected and concentrated to their final target concentrations before imaging by cryo-EM.

### ATPase activity assay

An NADH-coupled fluorimetric assay was used to measure ATPase activity (Scharschmidt et al., 1979). Mg^2+^-ATP was added to a mixture containing 0.046 to 0.092 µM hATP13A2, 50 mM Hepes, pH 7.5, 150 mM KCl, 30 mM MgCl_2_, 60 μg/mL pyruvate kinase, 32 μg/mL lactate dehydrogenase, 4 mM phosphoenolpyruvate, and 300 μM NADH. Consumption of NADH was measured by monitoring the fluorescence at *λ*_ex_=340 nm and *λ*_em_=445 nm using a SpectraMax Gemini microplate reader (Molecular Devices). Rates of ATP hydrolysis were calculated by converting fluorescence loss to nmol NADH per minute using known standards of NADH. Data were then fit by nonlinear regression to the Michaelis-Menten equation to calculate K_M_ and V_max_ values using GraphPad Prism. In lipid dependence experiments DPPA or PA were each supplemented at 0.1 mg/mL. Inhibition experiments were performed with the additional supplementation of: 1) 3 mM AMPPNP; 2) 5 mM MgCl_2_ and 10 mM NaF; or 3) 2 mM BeSO_4_ and 10 mM NaF.

### EM data acquisition

Purified hATP13A2 was concentrated to ∼5.0 mg/mL and supplemented with the following additives: i) 5.0 mM MgCl_2_, 2.0 mM AMPPNP; ii) 0.1 mM spermine, 5.0 mM MgCl_2_, 2.0 mM BeSO_4_, 10.0 mM NaF; or iii) 0.1 mM spermine, 5.0 mM MgCl_2_, 10.0 mM NaF. The supplemented protein sample was incubated at room temperature for ∼3 hrs and passed through a 0.45 µm filter to remove debris. To prepare cryo-EM grids, 3.5 µL drops of supplemented and filtered hATP13A2 were applied to Quantifoil R1.2/1.3 400 mesh Au grids glow-discharged for 60 sec. The grids were blotted for 4 sec at 4°C and 100% humidity before being plunge-frozen in liquid ethane using a Vitrobot Mark IV (FEI). The grids were imaged using a Titan Krios G3 transmission electron microscope (FEI) operated at 300 kV and a Gatan BioQuantum energy filter slit width of 5 eV. Automated data collection was performed with SerialEM using the beam-image shift method (Cheng et al., 2018; Mastronarde, 2003; Schorb et al., 2019). A Gatan K3 Summit direct electron detector was used to record movies in super-resolution counting mode with a super-resolution pixel size of 0.415 Å. Movies were recorded for 3.0 to 3.5 s over 60 or 70 frames using a dose-rate of 15 electrons per pixel per second with a defocus range of 0.6 to 1.8 µm or 0.5 to 1.3 µm. The total cumulative doses were 60 to 76 electrons per Å^2^ (1.09 electrons per Å^2^ per frame).

### Image processing and map calculation

The same data processing strategy was used in the early stages of structure determination to obtain all three hATP13A2 3D reconstructions presented in this study (Figure S4A). Briefly, dose-fractionated super-resolution movies were 2×2 down-sampled by Fourier cropping to a final pixel size of 0.83 Å. The down-sampled movie frames were used for grid-based motion correction and dose-filtering with MotionCor2 (Zheng et al., 2017). CTF parameters were estimated from the corrected movie frames using CTFFIND4.1 (Rohou and Grigorieff, 2015). The entire dataset was then inspected to eliminate micrographs exhibiting imaging defects including excessive drift, cracked ice or defocus values exceeding the specified range. Micrographs with estimated CTF resolution worse than 5 Å were discarded. The remaining motion-corrected dose-filtered movie sums were subjected to automated particle picking in RELION3.1 using a Laplacian-of-Gaussian filter to obtain an initial particle set (Kimanius et al., 2016; Scheres, 2012, 2016; Zivanov et al., 2018). Particle images extracted from the motion-corrected dose-filtered micrographs as 384×384 pixel boxes were down-sampled to 96×96 pixel particle stacks in RELION. Iterative 2D classification in RELION using down-sampled particle stacks was performed to remove spurious images of ice, carbon support and other debris.

For the E1-AMPPNP dataset, 740,626 particles from the best 2D classes were re-extracted without down-sampling as 384×384 pixel boxes in RELION (Figure S4B). *Ab initio* 3D classification was performed in cryoSPARC (Punjani et al., 2017) with C1 symmetry (Figure S4A). The *ab initio* models obtained were used as initial models for heterogeneous refinement resulting in one “good” class with clear protein features. The remaining “junk” classes were uninterpretable. 515,611 particles corresponding to the “good” class were subjected to a round of homogeneous refinement, resulting in a 3.43 Å reconstruction. Non-uniform refinement was performed using the refined 3D model to obtain a 3.00 Å reconstruction (Punjani et al., 2020). Lastly, local refinement was performed using a mask that includes the micelle density, which resulted in a final reconstruction at FSC=0.143 resolution of 2.9 Å (Figure S4E). Local resolution estimates mapped onto the final 3D reconstruction is shown in Figure S4H.

For the SPM-E2-BeF_3_^−^ dataset, the data processing strategy used was similar to that of the E1-AMPPNP dataset. Briefly, 1,368,235 particles from the best 2D classes were re-extracted without down-sampling as 384×384 pixel boxes in RELION (Figure S4C). *Ab initio* 3D classification was performed in cryoSPARC v2 with C1 symmetry to obtain three *de novo* classes (Figure S4A). The *ab initio* models obtained were used as initial models for of heterogeneous refinement resulting in one good class with clear protein features. The “good” class was then submitted to a round of non-uniform refinement. The refined particles were then used for a second round of heterogeneous refinement with the non-uniform refined model and “junk” models from the previous round of heterogeneous refinement as starting models. The resulting “junk” particles were eliminated, and the remaining “good” particles were subjected to another around of non-uniform refinement. The process of heterogeneous refinement and non-uniform refinement was iterated until the resolution of the non-uniform refined model no longer improves. The final round of non-uniform refinement using 648,247 particles yielded a refined 3D model at 3.13 A resolution. Lastly, local refinement was performed using a mask that includes detergent micelle density, resulting in a final reconstruction at FSC=0.143 resolution of 2.7 Å (Figure S4F). Local resolution estimates mapped onto the final 3D reconstruction is shown in Figure S4I.

For the SPM-E2-MgF_4_^2−^ dataset, a data processing strategy similar to that used for the SPM-E2-BeF_3_^−^ dataset was applied. Briefly, 466,778 particles from the best 2D classes were re-extracted without down-sampling as 384×384 pixel boxes in RELION (Figure S4D). *Ab initio* 3D classification was performed in cryoSPARC v2 with C1 symmetry to obtain three *de novo* classes. Iterative heterogeneous refinement and non-uniform refinement was then performed as described for the SPM-E2-BeF_3_^−^ dataset. The final round of non-uniform refinement using 285,797 particles resulted in a refined 3D model at 3.22 A resolution. Lastly, local refinement was performed using a mask that includes detergent micelle density, resulting in a final reconstruction at FSC=0.143 resolution of 3.0 Å (Figure S4G). Local resolution estimates mapped onto the final 3D reconstruction is shown in Figure S4J.

### Model building and coordinate refinement

Model building was initially performed in the E1-AMPPNP map. The crystal structure of rabbit skeletal muscle SERCA (PDB: 3N8G) was used as a reference structure to generate the starting model for building hATP13A2. Briefly, the SERCA model was first mutated generate a polyalanine model using CHAINSAW (Stein, 2008) and manually divided into its transmembrane domain and individual cytosolic domains. The domains were then docked into EM density separately by rigid-body fitting using the fitmap function in UCSF Chimera (Pettersen et al., 2004) followed by manual rebuilding in Coot (Emsley et al., 2010). The EM density was of sufficient quality to allow manual rebuilding in Coot. B-factor sharpening was performed locally in Coot “on-the-fly” to optimize observable map features for model building. The membrane embedded region of the NTD and the CTD were manually built *de novo* in Coot. The *β*-sandwich structure in the NTD was built using the N-terminal extension of the cryo-EM structure of yeast P5A-ATPase Spf (PDB: 6XMQ) as a starting model. The NTD exhibits a lower local resolution compared to the rest of the reconstruction and is built partly as a polyalanine model (a.a. 45 to 86). Rotamers were modeled for residues with poor sidechain density in the rest of the structure. Automatic real space refinement of the model against a half map was performed using `phenix.real_space_refine` (Adams et al., 2010). Tight secondary structure and geometric restraints were used to minimize overfitting. Manual rebuilding in Coot was alternated with automated refinement in `phenix.real_space_refine`. A similar strategy was used to build the SPM-E2-BeF_3_^−^ and SPM-E2-MgF_4_^2−^ structures, except the refined E1-AMPPNP coordinates was used as the starting model.

FSC curves were calculated between refined models and the half-map used for refinement (FSC_work_) or the half-map kept for cross-validation (FSC_free_) (Figure S4K to M). Model quality was evaluated by MolProbity (Chen et al., 2010) (Table S1).

Local resolution was estimated using `blocres` with a box size of 20 (Heymann and Belnap, 2007). The occlusion chamber and lumen access channel were identified using HOLLOW (Ho and Gruswitz, 2008) and MOLE 2.5 (Sehnal et al., 2013). All structure figures were generated using UCSF ChimeraX (Goddard et al., 2018).

## Supplementary Figure Legends

**Figure S1.**
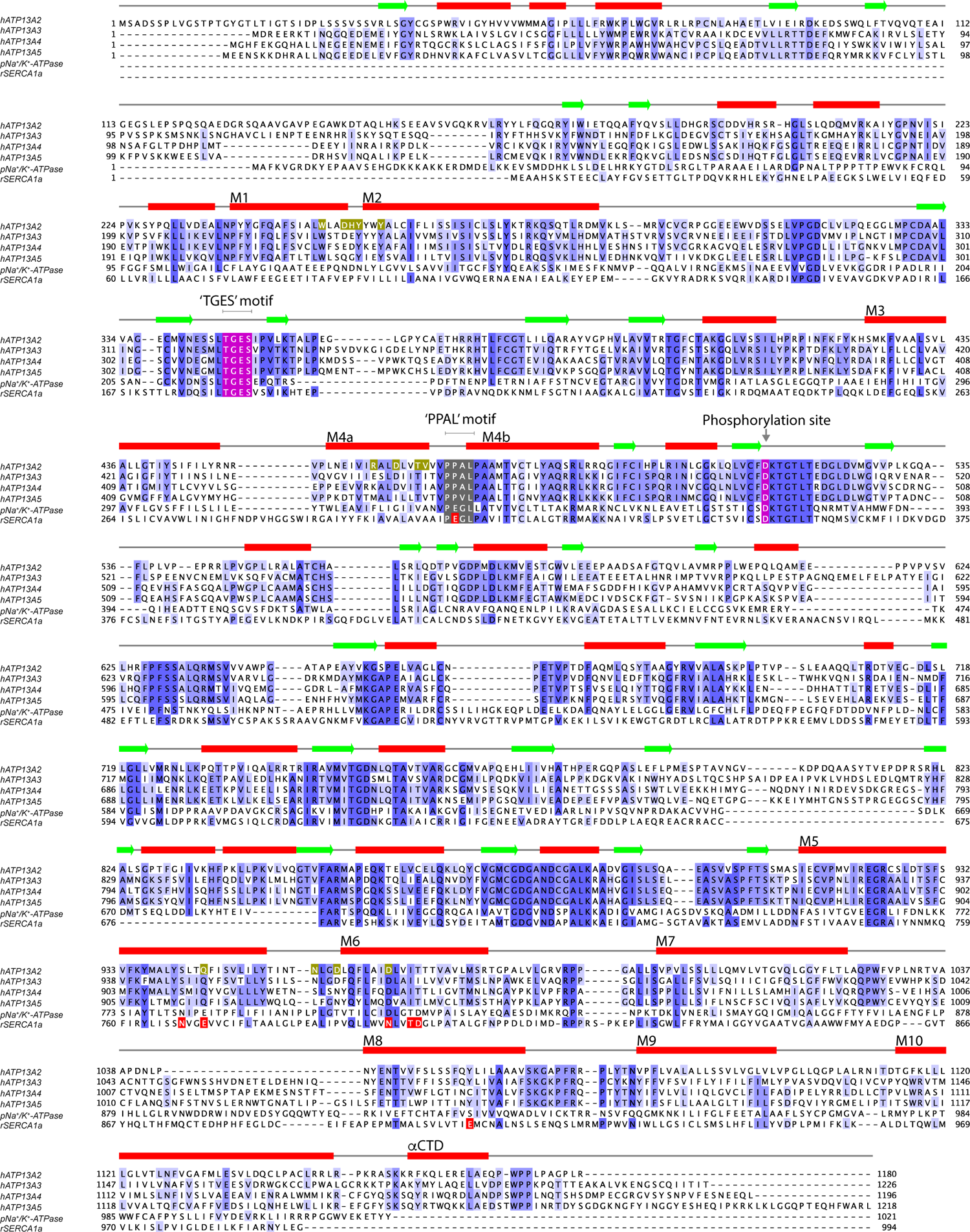
Sequence alignment of P5B-ATPases, related to. Figures 1**, 2, 3, 4, 5, 6, and 7.** A multiple sequence alignment of human P-type ATPases belonging to the P5B family. The protein sequence of rabbit SERCA and pig Na^+^/K^+^-ATPase are also included in the alignment. Sequence alignment was performed using ClustalW and adjusted manually. Shading in the sequence alignment indicate degree of conservation. Green arrows and red bars indicate *β*-strands and *α*-helices, respectively. Polyamine binding sites residues are highlighted by green boxes. Ca^2+^ binding site residues in SERCA are indicated by red boxes. The conserved TGES motif and the autophosphorylated aspartate residue are colored purple. The PPAL motif in the M4a/b-linker is indicated by gray shading.

**Figure S2.**
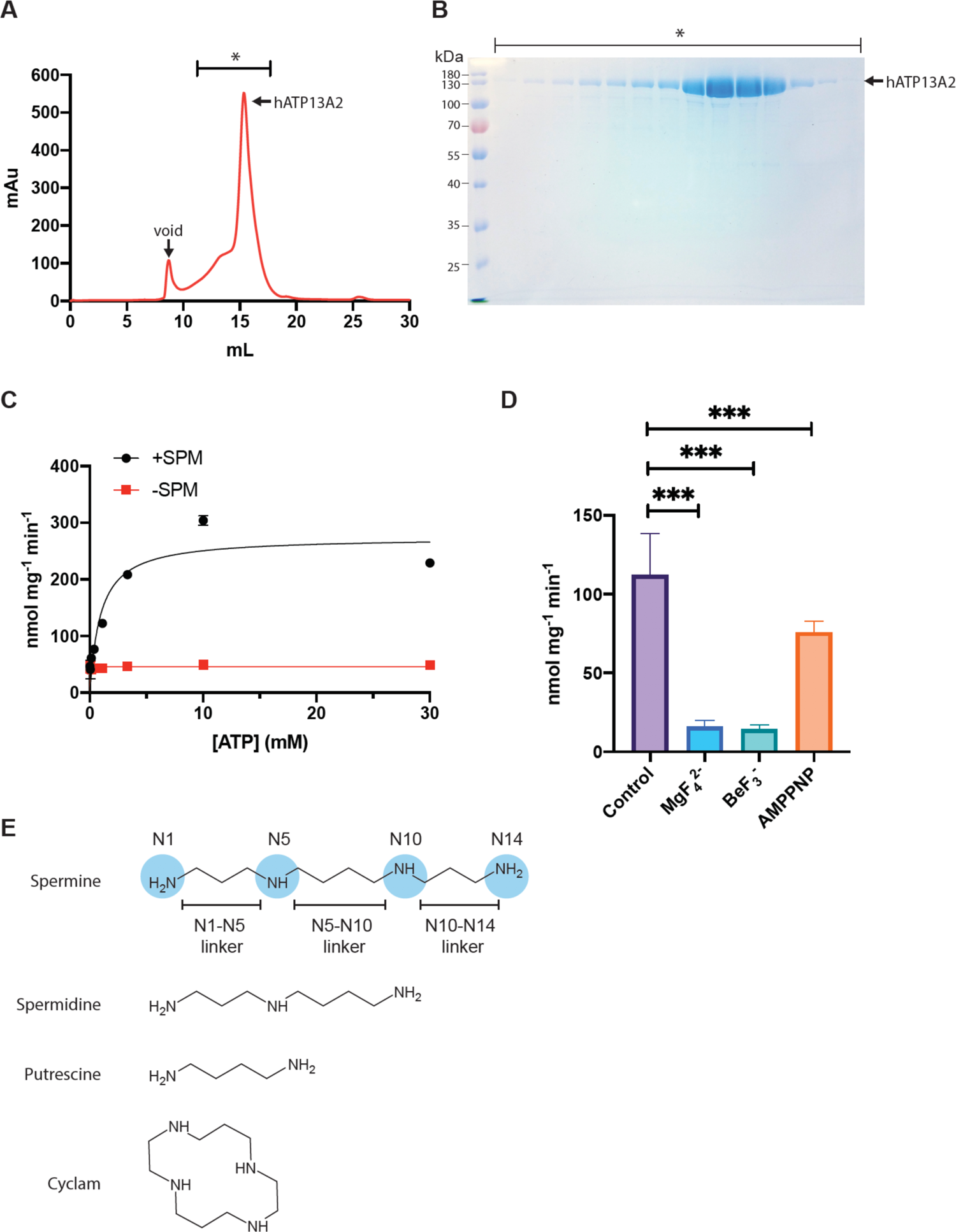
Purification and characterization of hATP13A2, related to. Figure 1. (A) Size-exclusion chromatography (SEC) profile of affinity purified hATP13A2. Asterisk indicates fractions analyzed by SDS-PAGE. (B) Coomassie stained SDS-PAGE gel of SEC purified hATP13A2. Asterisk indicates fractions in (A) there were analyzed. Predicted MW_hATP13A2_≈128.8 kDa. (C) Spermine-dependent ATPase activity of hATP13A2. ATPase activity was measured using purified hATP13A2 at 6 μg/mL in the presence or absence of 100 μM spermine. Data points represent mean ± SEM (n=3). The apparent K_M,ATP_ is 0.946 mM. (D) hATP13A2 ATPase activity in the presence of inhibitors. hATP13A2 is sensitive to inhibition by the phosphomimetics MgF_4_^2−^, BeF_3_^−^ or and the ATP analog AMPPNP. ATPase inhibition assays were performed in the presence of 10 μM spermine and 370 μM ATP. Data points represent mean ± SEM (n=3, ***P<0.001). One-way ANOVA analysis was performed. (E) Chemical structures of spermine, spermidine, putrescine and cyclam.

**Figure S3.**
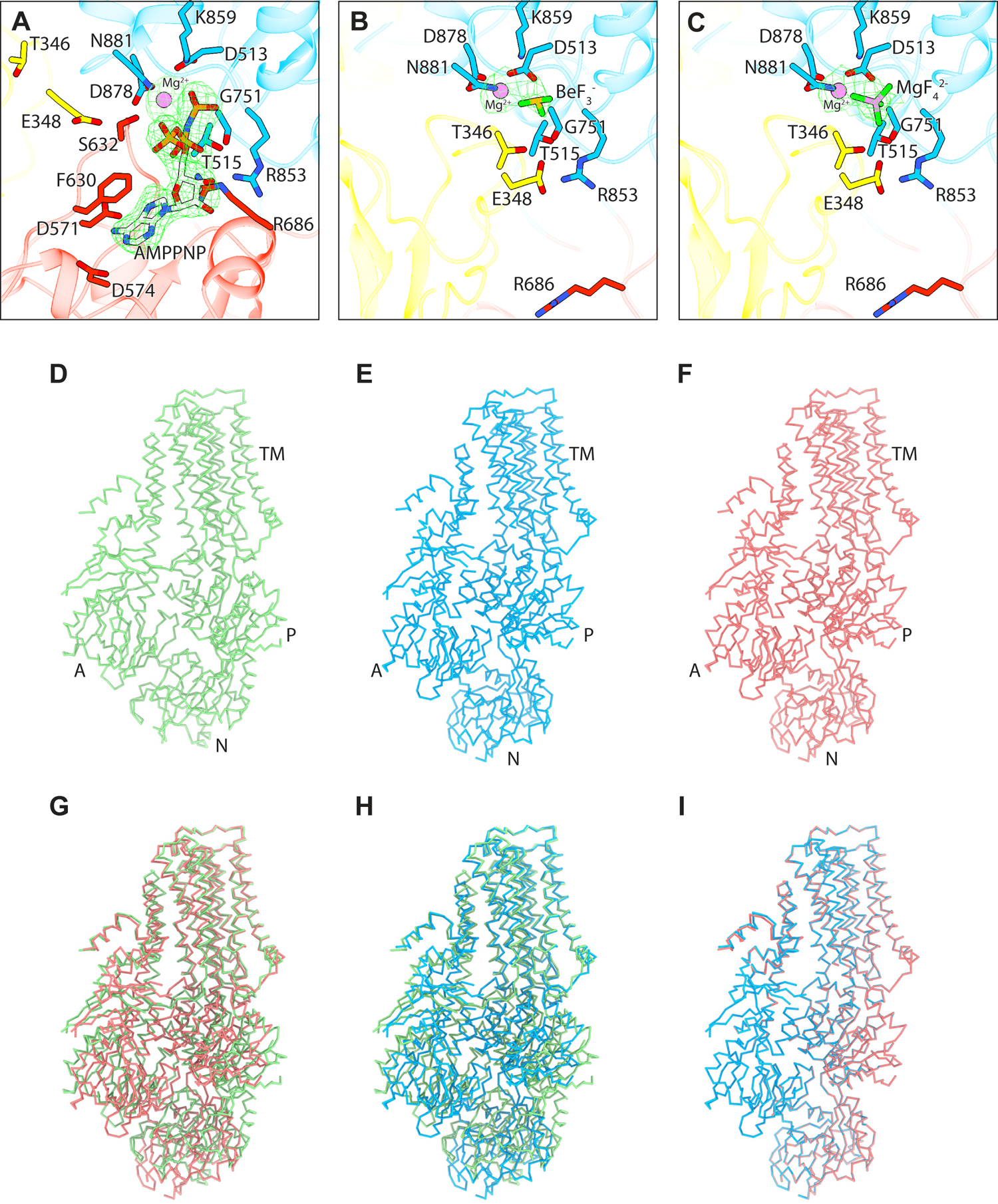
Comparison of hATP13A2 structures, related to. Figures 1**, 2, and 7.** (A to C) Zoomed in views of the ATPase active site in the E1-AMPPNP, SPM-E2-BeF_3_^−^ structure and SPM-E2-MgF_4_^2−^ structures, respectively. Coloring as in Figure 1D. Residues involved in autophosphorylation and dephosphorylation of Asp531 are shown. Cryo-EM densities corresponding to Mg^2+^-AMPPNP, Mg^2+^-BeF_3_^−^ and Mg^2+^-MgF_4_^2−^ are shown as green meshes. (D to F) *α*-carbon traces of the E1-AMPPNP (green), SPM-E2-BeF_3_^−^ (blue) and SPM-E2-MgF_4_^2−^ (red) structures, respectively. (G to I) Pairwise superpositions of hATP13A2 structures. Coloring as in D to F. (G) E1-AMPPNP vs SPM-E2-MgF_4_^2−^. (H) E1-AMPPNP vs SPM-E2-BeF_3_^−^. (I) SPM-E2-BeF_3_^−^ vs SPM-E2-MgF_4_^2−^.

**Figure S4.**
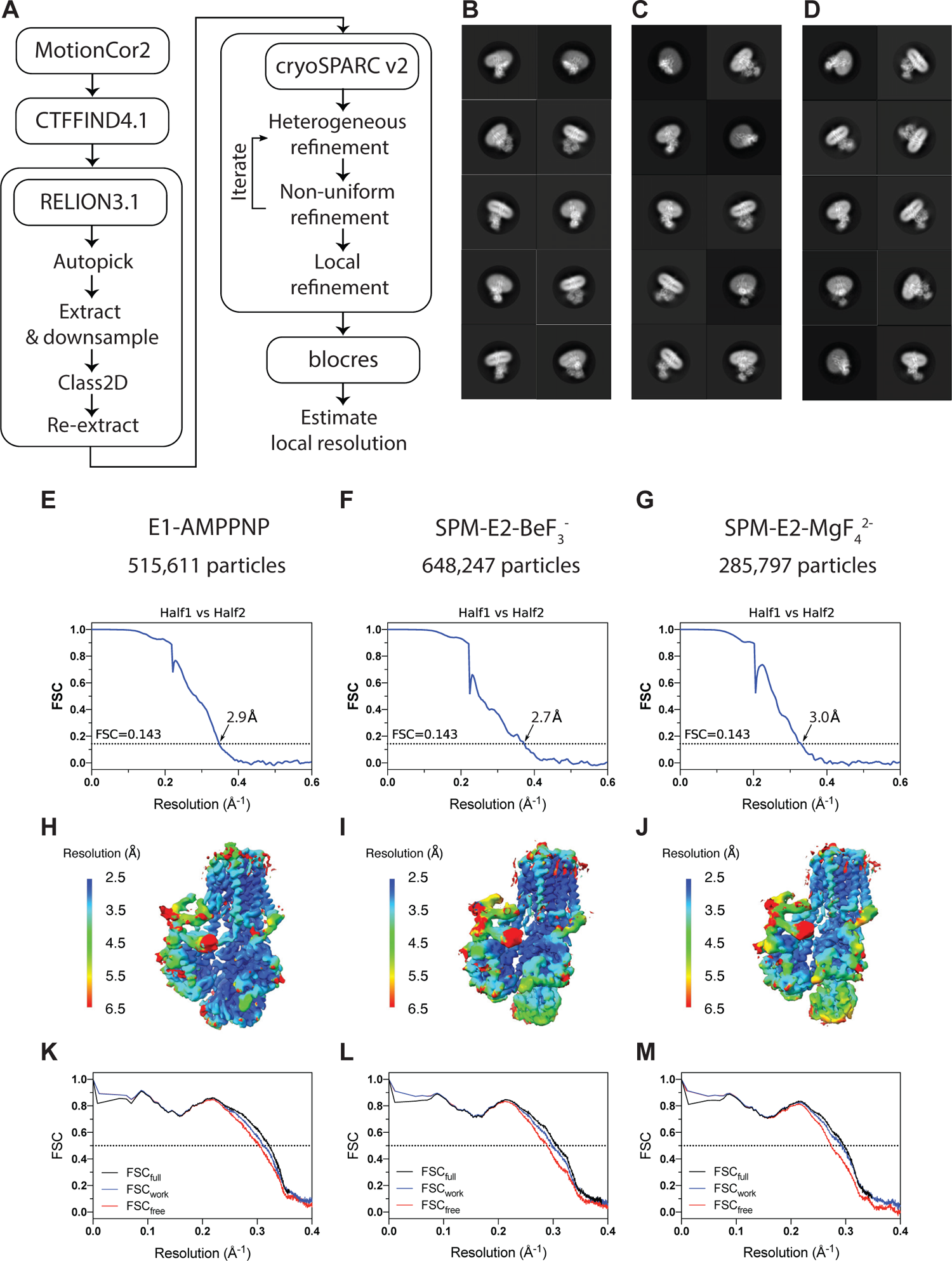
Image processing and resolution estimates, related to. Figures 1, 2, 3, and 4. (A) Outline of data processing strategy used for all datasets. (B to D) Representative 2D classes from the three datasets: E1-AMPPNP (B), SPM-E2-BeF_3_^−^ (C) and SPM-E2-MgF_4_^2−^ (D). (E to M) Estimates of global and local resolutions of the 3D reconstructions and the cross-validation of atomic models. (E, H and K) E1-AMPPNP structure. (F, I and L) SPM-E2-BeF_3_^−^ structure. (G, J and M) SPM-E2-MgF_4_^2−^ structure. (E to G) FSC curves calculated between independently refined half maps. (H to J) Local resolution maps. (K to M) Cross-validation FSC curves of the model vs. full map (FSC_full_), model vs. half map used for structure refinement (FSC_work_) and model vs half map excluded from structure refinement (FSC_free_).

**Figure S5.**
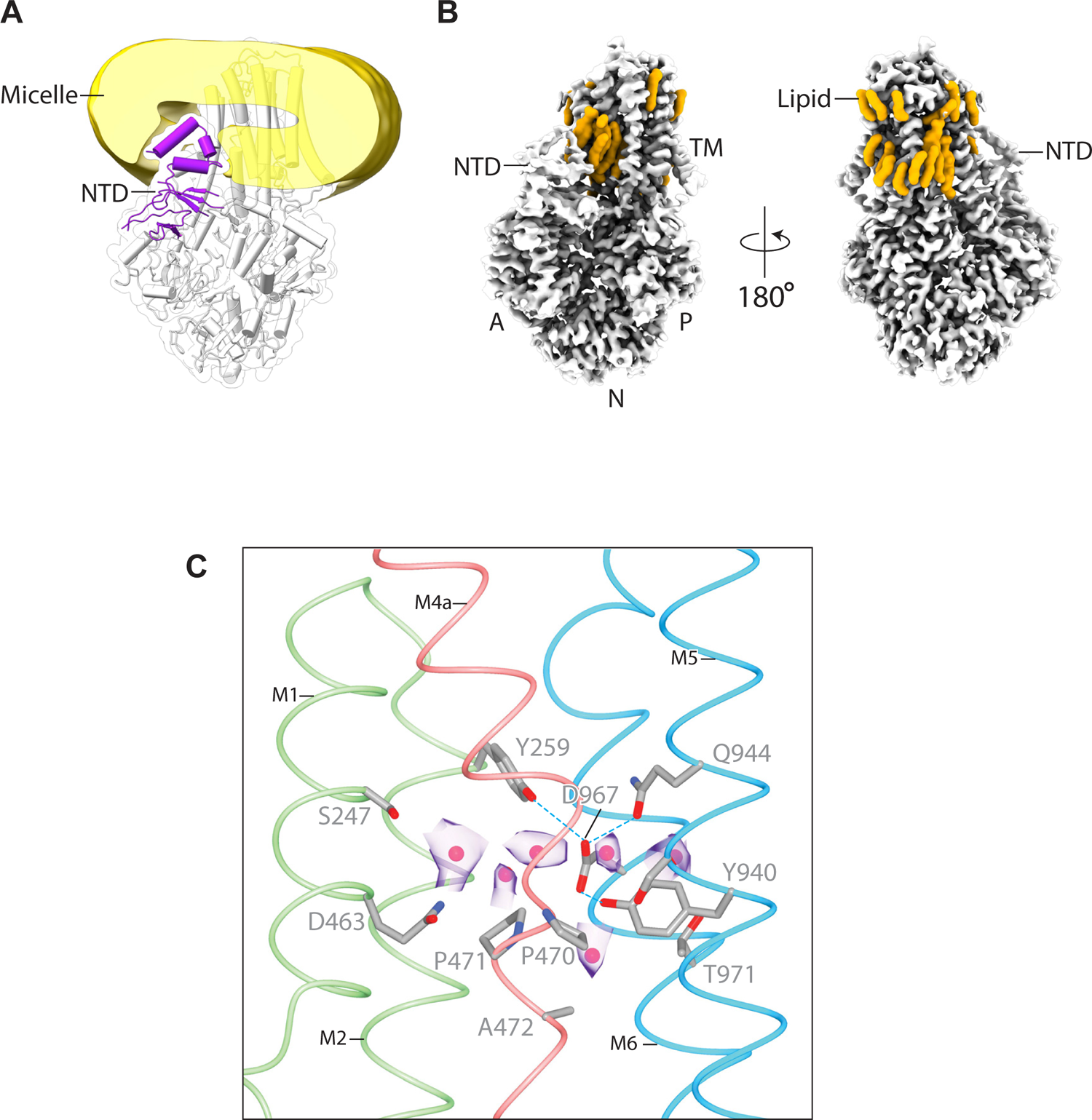
The lipid embedded NTD and the occlusion chamber in the E1-AMPPNP structure, related to. Figures 2**, and 3.** (A) The micelle density of the E1-AMPPNP structure. Cryo-EM density corresponding to the detergent micelle is low-pass filtered to 20 Å and shown along with the atomic-model of the E1-AMPPNP structure displayed in cartoon representation. The NTD is colored purple. The “spade” portion of the NTD is embedded in what can be considered the “inner leaflet” of the detergent micelle. (B) Lipid bilayer organization surrounding hATP13A2. Here lipid or detergent molecules bound to the periphery of hATP13A2’s transmembrane domain are shown in orange to delineate the lipid bilayer environment close to the protein. Cryo-EM density of the E1-AMPPNP structure is shown in gray. The insertion of the NTD in the inner leaflet can be readily appreciated. (C) Detailed view of the occlusion chamber in the E1-AMPPNP structure. Sidechains lining the occlusion chamber in the E1-AMPPNP structure are shown in ball-and-stick representation and colored in gray. Water molecules are shown as red spheres. Helices are colored as in Figure 3B. Cryo-EM density corresponding to occluded solvent is shown as transparent purple surfaces.

**Figure S6.**
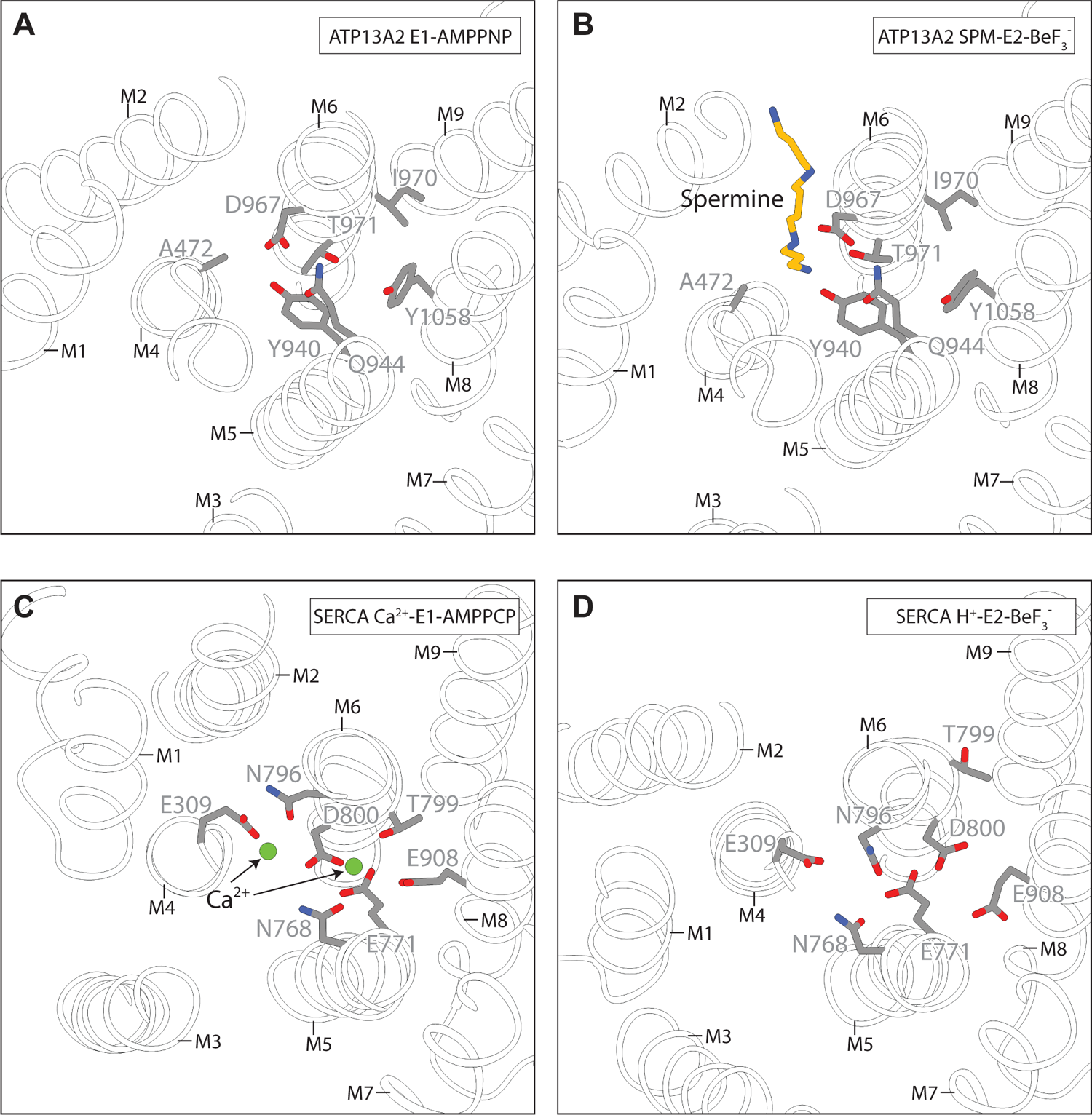
Comparison of SERCA and hATP13A2 structures, related to Figure, related to. Figures 3, 4, and 7. (A to D) Sidechains involved in Ca^2+^ coordination and proton binding in SERCA and the analogous residues in hATP13A2 are shown in ball-and-stick representation and colored gray. Ca^2+^ ions are shown as green spheres. Spermine is shown in ball-and-stick representation and colored yellow. Structures were aligned over M7 to M10. (A) hATP13A2 E1-AMPPNP. (B) hATP13A2 SPM-E2-BeF_3_^−^. (C) SERCA Ca^2+^-E1-AMPPCP (PDB ID 3N8G). (D) SERCA H^+^-E2-BeF_3_^−^ (PDB ID 3B9B)

**Figure S7.**
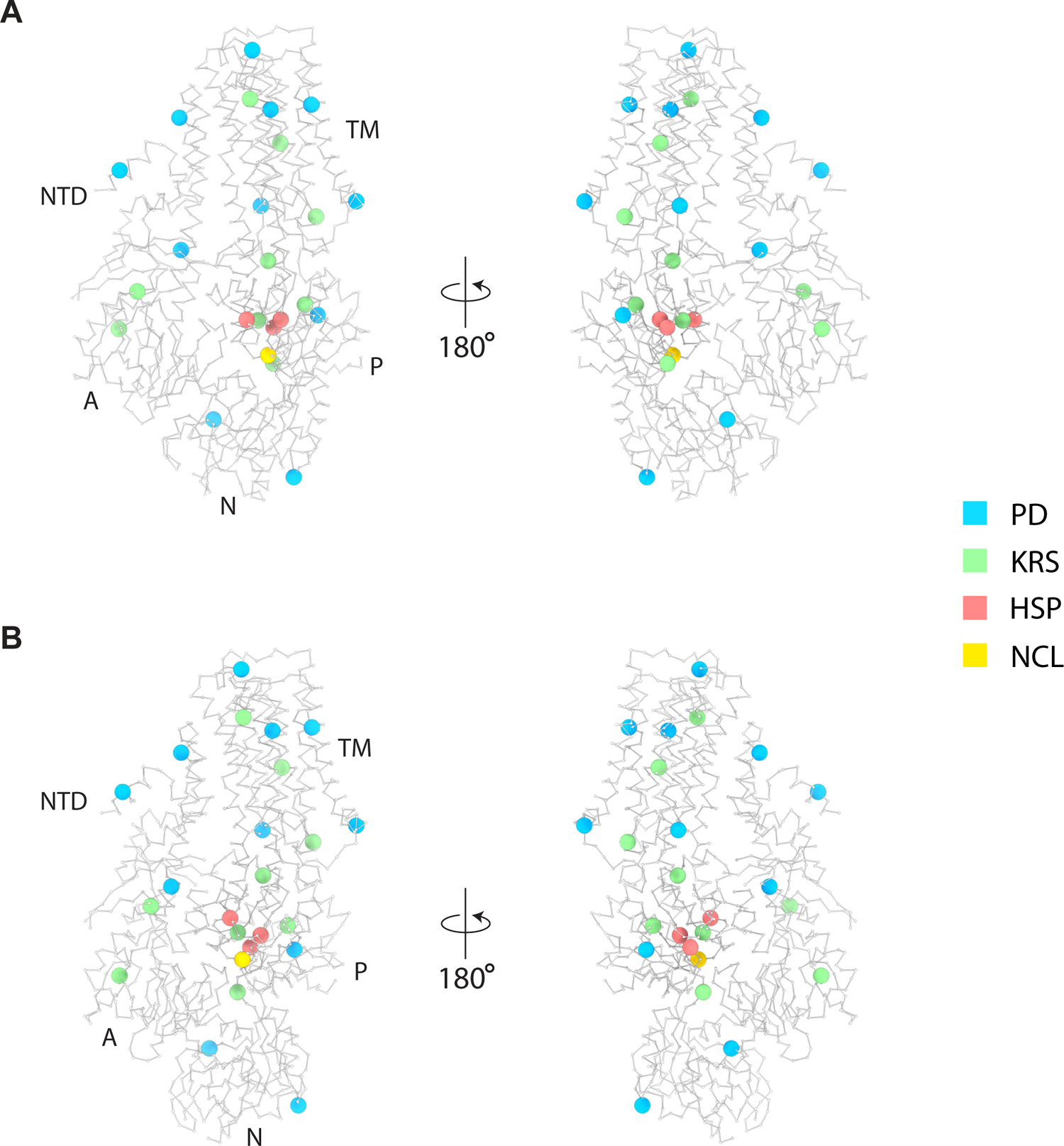
Disease-associated mutations in human ATP13A2, related to. Figure 7. (A and B) Mapping of ATP13A2 residues mutated in human diseases to hATP13A2 structures presented in this study. Gray *α*-carbon traces of hATP13A2 models are shown. *α*-carbon atoms of mutated residues are rendered as colored spheres according to the indicated coloring scheme. (A) hATP13A2 E1-AMPPNP. (B) hATP13A2 SPM-E2-BeF_3_^−^. PD – Early-onset Parkinson’s disease, KRS – Kufor-Rakeb syndrome, HSP – hereditary spastic paraplegia, NCL – neuronal ceroid lipofuscinosis.

**Table S1.** Data collection and refinement statistics, related to **Figures 1, 2**, 3, 4, 5, 6, and 7

|  | E1-AMPPNP | SPM-E2-BeF<sub>3</sub><sup>-</sup> | SPM-E2-MgF<sub>4</sub><sup>2-</sup> |
| --- | --- | --- | --- |
| Data acquisition |  |  |  |
| Microscope | Titan Krios | Titan Krios | Titan Krios |
| Voltage (kV) | 300 | 300 | 300 |
| Camera | Gatan K3 Summit | Gatan K3 Summit | Gatan K3 Summit |
| Camera mode | Super-resolution | Super-resolution | Super-resolution |
| Defocus range (μm) | 0.8 to 1.6 | 0.3 to 1.3 | 0.3 to 1.3 |
| Pixel size (Å) | 0.830 (super- | 0.830 (super- | 0.830 (super- |
| Movies | 7,969 | 18,231 | 7,336 |
| Frames/movie | 70 | 60 | 60 |
| Total electron dose | 76 | 65 | 65 |
| Exposure time (s) | 3.5 | 3.0 | 3.0 |
| Dose rate (e<sup>-</sup>/pixel/s) | 15 | 15 | 15 |
| Reconstruction |  |  |  |
| Software | cryoSPARC v2 | cryoSPARC v2 | cryoSPARC v2 |
| Symmetry | C1 | C1 | C1 |
| Initial Particle number | 740,626 | 1,368,235 | 466,778 |
| Final Particle number | 515,611 | 648,247 | 285,797 |
| Resolution (masked, Å) | 2.9 | 2.7 | 3.0 |
| Model statistics |  |  |  |
| Residues built | 1046/1180 | 1000/1180 | 1000/1180 |
| Map CC (masked) | 0.78 | 0.76 | 0.73 |
| Cβ outliers | 0 | 0 | 0 |
| Molprobity score | 1.66 | 1.58 | 1.60 |
| Clash score | 4.19 | 4.19 | 4.45 |
| Rama-Z score (whole) | 0.36 | 0.04 | 0.03 |
| Ramachandran |  |  |  |
| Favored (%) | 92.73 | 94.47 | 94.88 |
| Allowed (%) | 7.27 | 5.53 | 5.12 |
| Outliers (%) | 0.00 | 0.00 | 0.00 |
| RMS deviations |  |  |  |
| Bond length | 0.004 | 0.004 | 0.006 |
| Bond angles | 0.771 | 0.882 | 1.144 |

**Video S1. Allosteric mechanism for gating the lumen access channel**

## Notes

### Competing Interest Statement

The authors have declared no competing interest.

